# Acute chromatin decompaction stiffens the nucleus as revealed by nanopillar-induced nuclear deformation in cells

**DOI:** 10.1101/2025.03.06.641794

**Authors:** Aninda Mitra, Marie FA Cutiongco, Romina Burla, Yongpeng Zeng, Na Qin, Mengya Kong, Benjamin Vinod, Mui Hoon Nai, Barbara Hübner, Alexander Ludwig, Chwee Teck Lim, GV Shivashankar, Isabella Saggio, Wenting Zhao

## Abstract

Chromatin architecture is critical in determining nuclear mechanics. Most studies focus on the mechanical rigidity conferred by chromatin compaction from densely packed heterochromatin, but less is known on how transient changes in chromatin decompaction state impinge on nucleus stiffness. Here, we used an array of vertically aligned nanopillars to study nuclear deformability *in situ* after chromatin decompaction in cells. The nucleus significantly stiffened within 4 hours of chromatin decompaction but softened at longer timescales. This acute nucleus stiffening was predominantly underlied by an increase in nucleus volume, nuclear import and partially enhanced by lamin protein recruitment to the nuclear periphery. The coupling between nucleus stiffening and acute chromatin decompaction was observed in cancer cell lines with lower malignancy (e.g. MCF7, PEO1, A549) but weakened in those with higher metastatic potential (e.g. MDA-MB-231, HEYA8, HT1080), which was found to be associated with the capacity to efficiently compact heterochromatin into foci that sustains nucleus deformability required for confined migration. Our work signals how a rapid chromatin remodeling is a physiologically relevant pathway to modulate nucleus mechanics and cell migration behavior.

**STATEMENT OF SIGNIFICANCE:** Many cell processes such as wound healing, immune activation and DNA damage repair require a decompact and accessible chromatin structure. Whether such short-term remodeling of the chromatin impacts nucleus mechanics and function is poorly defined. Using nanopillars that allow interrogation of nucleus rigidity within intact cells, we showed that contrary to conventional knowledge the nucleus becomes less deformable and more rigid when chromatin is acutely decompacted due to enhanced nuclear import and swelling of the nucleus. In cancer cells, the coupling of transient chromatin decompaction to nucleus rigidity is weakened and appears to be countered by heterochromatin formation and compaction. We demonstrate here how short-term chromatin remodeling can impact nucleus and cellular properties in a time-dependent and non-genetic manner.

## INTRODUCTION

Aside from its primary function in determining gene expression, chromatin is now known to contribute to the physical properties of the nucleus^1^ through its dynamic and variable compaction state^2–4^. Studies mainly attribute nuclear mechanical rigidity to heterochromatin, which is densely packed and consists of transcriptionally-repressed genes and methylated histones, and attaches to and supports the nuclear envelope^5–7^. In fact, loss of heterochromatin is sufficient to soften the nucleus^4,7,8^. Conversely, studies show that chromatin decompaction through genetic modification reduces the rigidity and sturdiness of the nucleus^5^. Small molecules for histone deacetylation (HDAC) inhibition-based chromatin decompaction (such as trichostatin A and valproic acid) similarly exhibit a softening effect on the nucleus when used long-term^3^. Independent of the nuclear lamina, chromatin acts as a primary structure determining nuclear integrity, and a change in its condensation state influences nucleus stiffness^3^ and impinges on cellular behavior such as cell migration^9^. What remains less explored is whether cells can transiently alter nuclear rigidity and cellular behavior via chromatin remodeling. In particular, the consequences of acute chromatin decompaction arising from cell processes such as T cell activation^10^ and wound healing^11^ on nuclear mechanics are underappreciated.

Vertically aligned, high-aspect ratio nanopillars are useful tools that can provide controllable indentation and guidance to cellular membranes at the nanoscale. Nanopillars have thus been used in a wide range of applications to study and engineer the nuclear-cytoskeleton mechanotransduction pathway to alter stem cell fate^12,13^, and enhance transfection efficiency and delivery of biomolecules^12,14^. Nanopillars also have the unique capability of probing deeply into the cell, deforming both the cellular^15–18^ and nuclear membranes^19–21^. The deformation of the nucleus by nanopillars is a read-out of nucleus stiffness, where softer nuclei exhibit a tighter contouring of the nuclear membrane against the nanopillars and a deeper indentation of nanopillars into the nucleus^19^. This readily measures nucleus stiffness *in situ* and in live, intact cells and is ideal for exploring the effect of chromatin on nuclear mechanics and cell behaviour.

Here, we used nanopillars to examine how a temporal change in chromatin decompaction alters nucleus mechanical properties. Pharmacological and genetic tools were utilized to induce chromatin decompaction at different time scales, upon which the deformability of nuclei on nanopillars was measured. In contrast to prevailing knowledge, chromatin decompaction acutely and counter-intuitively led to a rigidified nucleus. To assess the underlying mechanism, we then tested the contribution of intranuclear pressure and the nuclear lamina to nucleus stiffening after chromatin decompaction. We then tested breast, ovarian and lung cancer cell lines with different malignancy levels, where we found the coupling between nucleus stiffening and transient chromatin decompaction to be dependent on malignancy and capacity for heterochromatin formation.

## RESULTS and DISCUSSION

### Chromatin decompaction causes increase in nucleus stiffness

We used the histone deacetylase (HDAC) inhibitor Trichostatin A (TSA) to reversibly and globally increase chromatin acetylation and decompaction, reduce condensation of chromatin, and increase chromatin spatial homogeneity^22–24^. Hela cells on unpatterned coverslip showed increased euchromatic H3K9ac intensity, nuclear area, and volume after treatment with TSA (250 ng/mL, 4 hours) compared to control (Supplementary Figure 1). Transmission electron microscopy (Supplementary Figure 2) confirmed chromatin decompaction after TSA treatment, revealing a reduction in heterochromatic regions along the nuclear envelope and around nucleoli, and more decondensed chromatin in most other regions of the nucleus compared to control cells.

We then used vertically aligned nanopillars to probe *in situ* the nanoscale mechanical architecture of the nucleus, whose stiffness determines the depth of deformation induced by nanopillars^19^ (Fig. 1a). Hela cells on nanopillars exhibited global increase in H3K9ac intensity after 4 hours of TSA treatment (Fig. 1b). Visualization of the nuclear envelope using antibodies against Lamin A reveal ring-like structures colocalizing with nanopillars in control nuclei, with less prominent patterns featured in TSA-treated nuclei (Fig. 1b). This recapitulates previous reports where the nuclear envelope molds itself against the contour of nanopillars and reveals ring patterns^19,25^, with overexpression of Lamin A/C and rigidification of the nucleus leading to less visible patterns^19^. A cross-section of the ring patterns induced by nanopillars yielded images resembling pillars (Fig. 1c). By following the changes in the intensity profile of Lamin A imaged along the entire volume of the nucleus, we calculated the nanopillar-induced deformation depth to infer nucleus deformability (Supplementary Figure 3). We observed that 4 hours of TSA treatment significantly reduced deformation depth compared to control nuclei, thus signifying an increase in nucleus stiffness after acute chromatin decompaction (Fig. 1d). To support our measurements using diffraction-limited confocal microscopy, we utilized expansion microscopy study with nanoscale resolution the changes in nuclear deformability after TSA treatment. Images of the nuclei stained against Lamin A and nanopillars coated with fluorescently labelled gelatin clearly show the nanopillars indenting the nuclear envelope more deeply inwards in control compared to TSA-treated nuclei (Fig. 1e-f, Supplementary Figure 4). Since expansion microscopy provides nanometer resolution, we were able to directly measure nanopillar-induced deformation depth from cross-sectional views of Lamin A centered on nanopillars. Deformation depth measurements from expansion microscopy imaging support our observations using conventional microscopy, that nuclei treated with TSA for 4 hours become harder to deform compared to control nuclei (Fig. 1g, Supplementary Figure 5).

**Figure 1.**
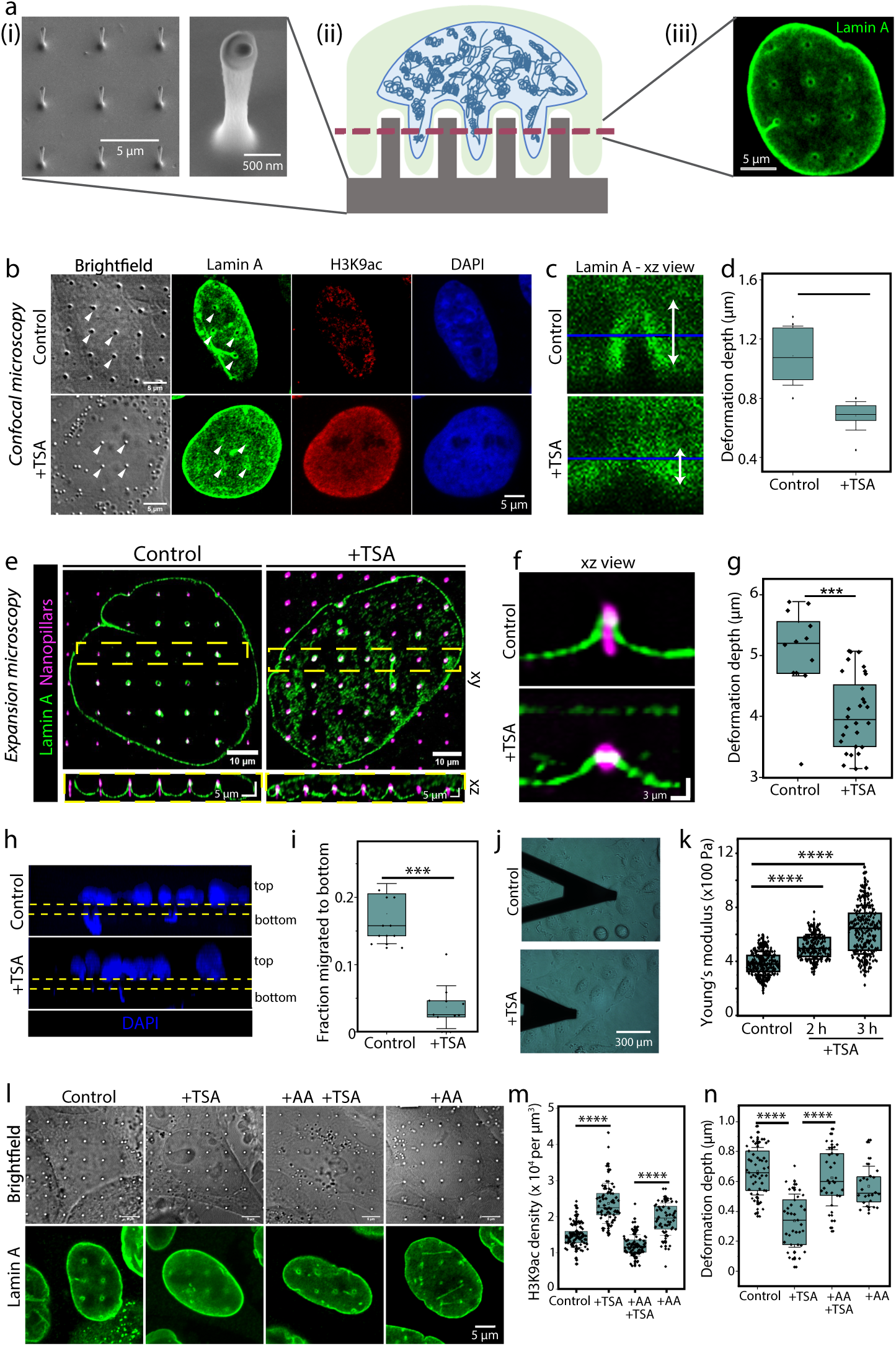
Chromatin decompaction leads to stiffening of the nucleus. (a) Schematic of vertically aligned nanopillars used to deform the nuclei. i, Representative scanning electron micrograph (SEM) of vertically aligned nanopillar array. ii, Schematic representation showing cells attached to nanopillar array (cross-section). iii, Example image of nucleus stained for Lamin A after attachment to nanopillars. (b) Representative images of H3K9ac and Lamin A after treatment with Trichostatin A (TSA) for 4 hours in Hela cells on nanopillars. Hela without treatment were used as a control. White arrowheads show the location of pillars in brightfield and Lamin A images. (c) Representative cross-section of nanopillar-induced deformation of the nucleus (visualized by Lamin A) under untreated and TSA-treated regimes. (d) Nanopillar-induced deformation depth measured from untreated and TSA-treated nuclei. (e) Representative control and TSA-treated nuclei imaged for Lamin A and fluorescence-coated nanopillars under expansion microscopy. Pillar array highlighted in the yellow dashed box are shown in xz (cross sectional) view. (f) Representative xz view from expansion microscopy of Lamin A deformation on fluorescence-coated nanopillars in control and untreated nuclei. (g) Nanopillar-induced deformation depth measured from untreated and TSA-treated nuclei imaged under expansion microscopy. (h) Representative image of cells migrating through transwell membrane (5 µm pore size) under TSA and Control conditions. Yellow dashed band indicates location of transwell membrane and blue denotes the nucleus stained against DAPI. (i) Fraction of nuclei that migrate to the bottom of the transwell for both Control and TSA-treated conditions. (j) Representative image of atomic force microscopy (AFM) cantilever on cells. (k) Young’s modulus of nuclei measured from Control and TSA-treated cells. (l) Representative images of Hela nuclei on nanopillars under untreated, TSA, Anacardic acid (AA) or both TSA and AA treatment conditions. (m) H3K9ac intensity per nucleus volume measured from nuclei under untreated, TSA, Anacardic acid (AA) or both TSA and AA treatment conditions. (n) Nanopillar-induced deformation depth measured from nuclei under untreated, TSA, Anacardic acid (AA) or both TSA and AA treatment conditions.

Transwell migration and measurement of nuclear elastic moduli using atomic force microscopy (AFM) support our measurements of nanopillar-induced nuclear deformability. Nucleus stiffness^26,27^ (and specifically stiffness of the nuclear lamina^28^) impedes 3D migration through a transwell, which was observed from the lower fraction of Hela cells that migrated to the bottom of the transwell under TSA treatment compared to control (Fig. 1h-i). AFM of individual nuclei correlated with these measurements, showing an increase in Young’s modulus after acute TSA treatment (Fig. 1j-k). Finally, we tested the reversibility of the nucleus stiffening effect induced by chromatin decompaction using the histone acetylase inhibitor anacardic acid (AA, Fig. 1l). Treatment of cells with AA alone did not yield significant changes in either H3K9ac density (Fig. 1m, Supplementary Figure 6) or nanopillar-induced deformation depth (Fig. 1n) compared to the control. Meanwhile, combined treatment of TSA and AA recovered both H3K9ac density and nanopillar-induced deformation depth from TSA back to control levels (Fig. 1m-n) signaling the dynamic and transient nature of chromatin decompaction effects on nuclear deformability.

### Stiffening of the nucleus upon chromatin decompaction is time dependent

The nucleus rigidification observed here contradicts prior reports on the softening effect of TSA (and chromatin decompaction in general) on the nucleus. One major difference between our work and previous work is the use of a shorter timepoint for TSA treatment, whilst phenomena were studied at longer (≥16 hours) timescales previously^3,24,26^. We then next examined the time dependent effect of chromatin decompaction on nuclear deformability (Fig. 2a). Hela cells on nanopillars were treated with TSA for different durations, then visualized using H3K9ac intensity and Lamin A (Fig. 2b). H3K9ac density peaked at 4 hours post-TSA treatment followed by a decline at 24-hour post-TSA treatment (Fig. 2c). Nanopillar-induced deformation depth dipped at 4 hours post-TSA treatment compared to control, then increased to a level higher than the control at 24 hours after TSA treatment (Fig. 2d). The change in nucleus deformability correlated with changes in nucleus volume (Fig. 2e), which also peaked at 4 hours post-TSA treatment. Interestingly, both H3K9ac intensity and nucleus volume started to decline at 6 hours post-TSA treatment (Supplementary Figure 7). AFM measurements of cells verified that the Young’s modulus of TSA-treated nuclei peaked at 4 hours then gradually started to recover after 24 hours (Fig. 2f). Tracking nanopillar-induced deformation in individual cells using live microscopy gave visual evidence at the single cell level of a steady decrease in deformation depth from 0 h to 4 h post-TSA treatment (Fig. 2h-I, Supplementary Video 1). We demonstrated here that chromatin decompaction affects nucleus mechanics in a time dependent manner, and that acute chromatin decompaction leads to nucleus stiffening. This contradicts conventional knowledge wherein long-term chromatin decompaction leads to increased mobility or flow of chromatin, loss of heterochromatin tethering to the nuclear envelope and decreased mechanical reinforcement of the nuclear lamina^7^. The transience of nucleus stiffening after acute chromatin decompaction suggests that this process may be a homeostatic mechanism to cope with the expansion of chromatin to ensure maintenance of nuclear form and function and could be valuable in processes that require a burst of chromatin decompaction such as in fate determination^29^, B-cell activation^30^, mechanosensing^31^ and DNA repair^32^ responses (which are all transcription-heavy activities).

**Figure 2.**
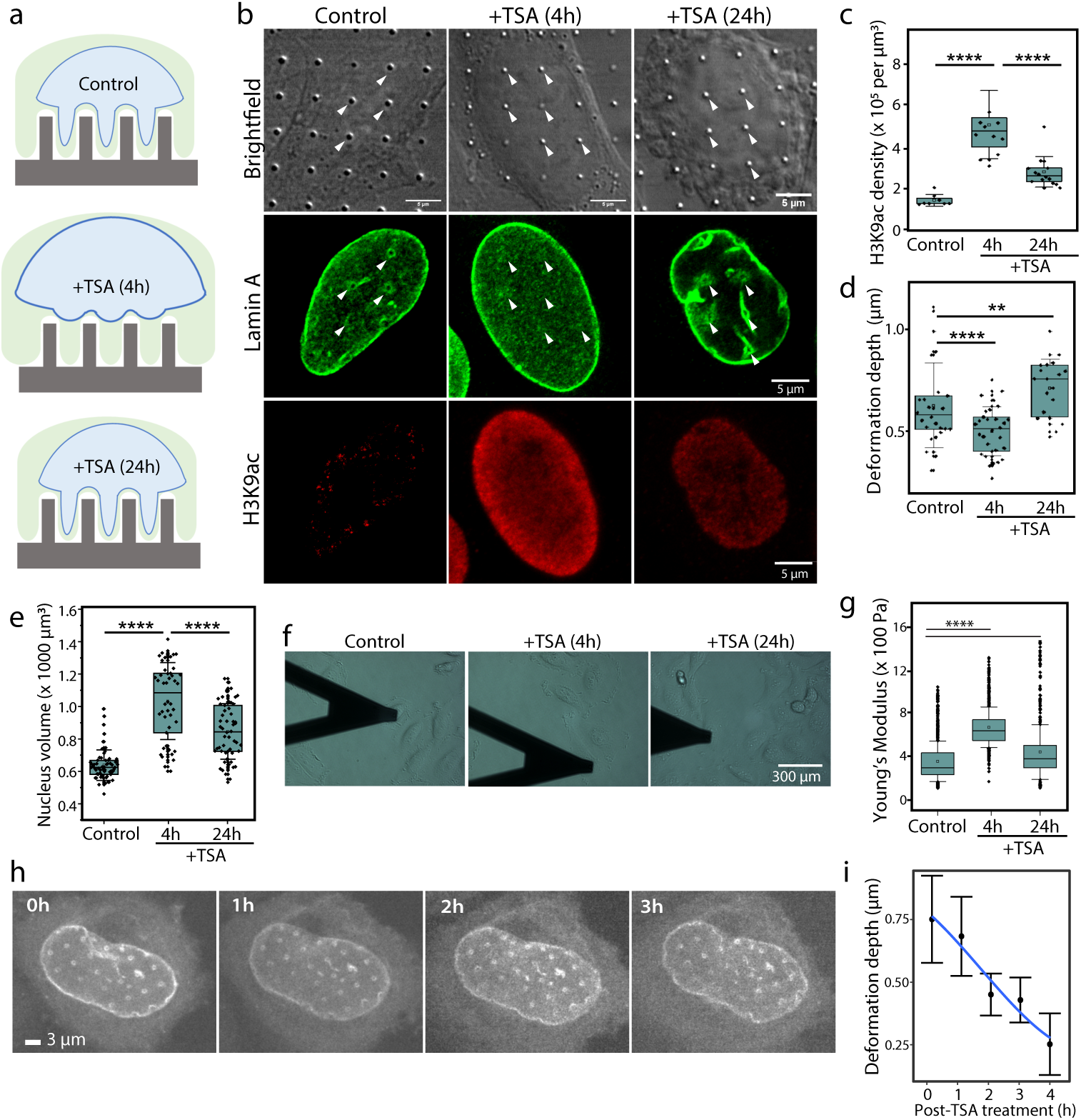
Chromatin decompaction and nucleus stiffening induced by TSA is time dependent. (a) Schematic of using vertically aligned nanopillar array to measure nanopillar deformation under acute (4h) and chronic (24h) TSA treatment. (b) Representative images of Lamin A and H3K9ac after acute and chronic TSA treatment of Hela cells on nanopillars. White arrowheads show the location of pillars in brightfield and Lamin A images. (c-e) Measures of (c) H3K9ac density, (d) nanopillar-induced deformation depth and (e) nucleus volume depends on the duration of TSA treatment. (f-g) Nuclei stiffness measured using AFM. (f) Representative AFM images of cells treated with TSA at different durations. (g) Young’s modulus of cells depends on the duration of TSA treatment. (h) Representative image of nuclei transfected with Lamin A-chromobody and tracked hourly using timelapse microscopy. (i) Nanopillar-induced deformation depth tracked from the same Hela cells over time. (8 cells tracked via live cell microscopy).

TSA substantially influences cell cycle progression and cell death such that it has been tested as an antitumor drug. Previous work showed that Hela cells required at least 12 hours of TSA treatment to see weak arrest at the G1 and G2/M phases using 100 ng/ml^33^ or a significant drop in cell proliferation rate using 500 ng/ml^34^ of TSA. Similarly, 12 hours of TSA treatment at 150 ng/ml failed to induce any remarkable change in cell cycle progression of MCF7 and MDA-MB-231^35^. At the 12 hour timepoint, chromatin decompaction-induced nucleus stiffening was no longer observed in Hela cells (Supplementary Figure 7). Additionally, the apoptotic effect of TSA has only been shown for treatment time of 24 hours or longer^35–38^. The literature signifies that at the 4-hour timepoint, the antiproliferative effects of TSA is negligible compared to the significant changes seen in nuclear shape, stiffness and deformability we report here.

### Stable chromatin decompaction similarly leads to acute stiffening of the nucleus

We next explored the effect of genetically depleting the protein AKTIP, which leads to stable chromatin decompaction^39^. Using siRNA against AKTIP (‘siAKTIP’) we effectively knocked down expression of AKTIP protein (Supplementary Figure 8), causing an increase in H3K9ac intensity and nucleus volume compared to an siRNA control (Fig. 3a-c). Similar to acute TSA treatment, siAKTIP nuclei exhibited reduced nanopillar-induced deformability compared to control nuclei (Fig. 3d-e). The decrease in nuclear deformability in siAKTIP nuclei was supported by higher Young’s modulus captured via AFM (Fig. 3f-g) and reduced transwell migration compared to siControl (Fig. 3h-i). Whereas 24-hour treatment with TSA led to a softer nucleus, chromatin decompaction through genetic modification of chromatin remodeling proteins causes an permanent increase in nucleus stiffness.

**Figure 3.**
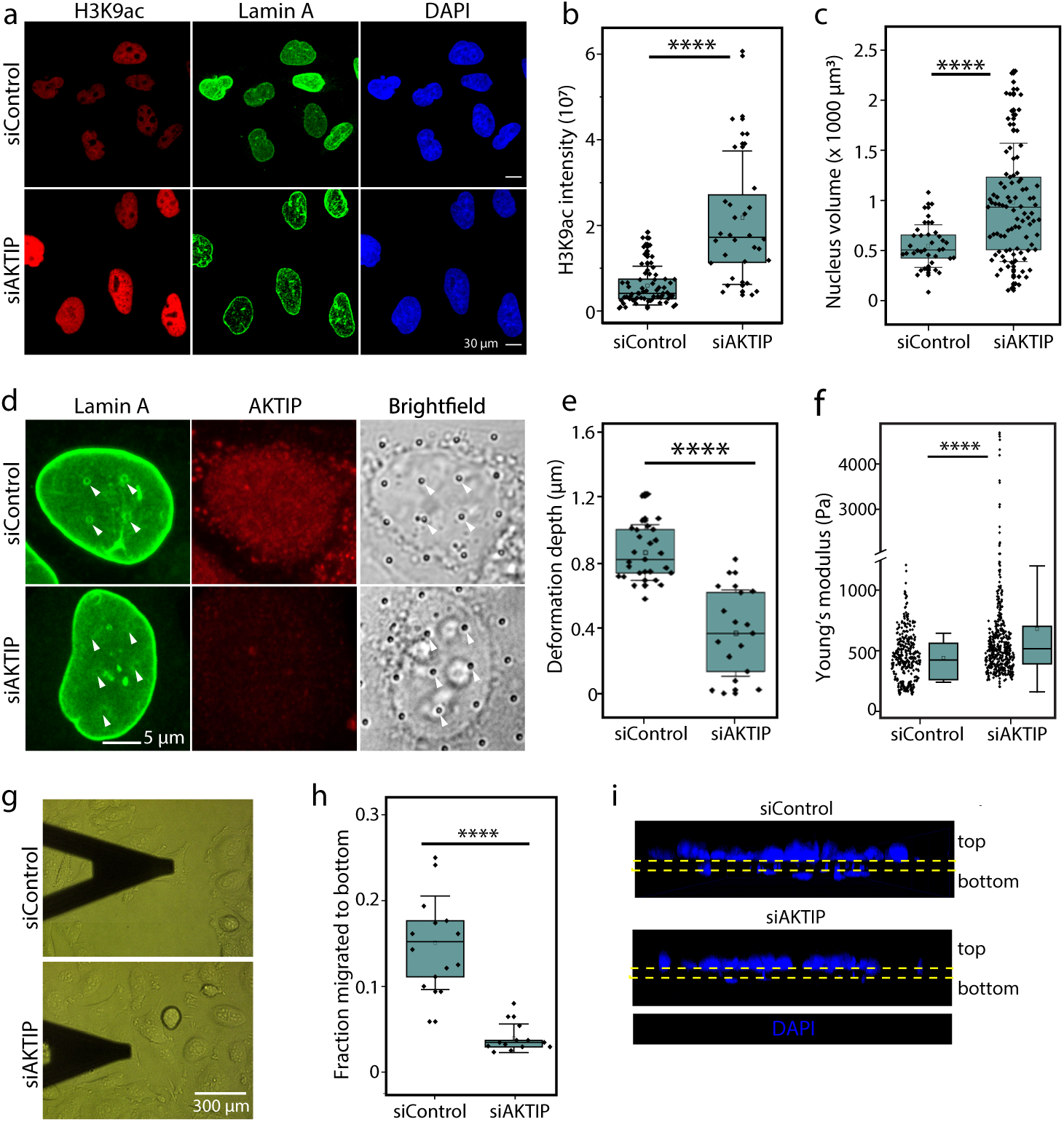
AKTIP-induced stable chromatin decompaction stiffening of the nucleus. (a) Representative images of Hela cells modified with siRNA against AKTIP (siAKTIP) and scramble control (siControl). Nuclei were stained against H3K9ac, Lamin A and DAPI. (b) H3K9ac intensity per nucleus measured from siControl and siAKTIP cells. (c) Nucleus volume measured from siControl and siAKTIP cells. (d) Representative images of Lamin A and AKTIP in siControl and siAKTIP cells cultured on nanopillars. White arrowheads show the location of pillars in brightfield and Lamin A images. (e) Nanopillar-induced deformation depth measured in siAKTIP and siControl cells. (f) Young’s modulus of siAKTIP and siControl nuclei using AFM. (g) Representative image of atomic force microscopy (AFM) cantilever on siAKTIP and siControl cells. (h) Fraction of cells migrated through transwell in siAKTIP and siControl cells. (i) Representative image of cells after migration through 5 µm pore size transwell. Yellow dashed band indicates location of transwell membrane and blue denotes the nucleus stained against DAPI.

### Nucleus stiffening induced by chromatin decompaction primarily involves nuclear import and the increased recruitment of lamins

There are two possible but opposing forces that underlie the increase in nucleus stiffness in response to acute chromatin decompaction: (1) an increase in nuclear lamina stiffness, purported to be the main mechanical component of the nuclear envelope; concurrently with or alternative to (2) an increase in intranuclear pressure from expansion of the chromatin (evidenced by an increase in nucleus volume).

The first hypothesis was supported by Western blotting showing elevated total protein levels of Lamins A, C and B1 after 1.5 hours TSA treatment compared to control (Supplementary Figure 9). Image analysis of nuclei treated with TSA for 4 hours revealed that both Lamin A and Lamin B1 intensity in the nucleus increased compared to control (Fig. 4a-b). In TSA treated cells, the change in total lamin level was accompanied by localization of Lamin A and Lamin B1 proteins to the nuclear periphery compared to the nucleoplasm (Fig. 4d-e). A similar trend was observed for siAKTIP versus siControl nuclei, where Lamin A and Lamin B1 were brighter at the nucleus periphery (Supplementary Figure 10). To verify whether the increased localization of lamin proteins at the nuclear periphery was solely responsible for nucleus stiffening under a decompacted chromatin state, we overexpressed lamin proteins to modulate nucleus stiffness. We hypothesized that combined acute TSA treatment and Lamin A overexpression would lead to a nucleus stiffened beyond what TSA treatment will induce (Fig. 4f). Acute TSA treatment of cells with GFP or GFP-Lamin A/C overexpression induced the characteristic response of increased nucleus volume (Fig. 4g) and reduced nanopillar-induced deformability (Fig. 4h) compared to control. Surprisingly, there was no observable difference in these parameters when comparing GFP and GFP-Lamin A/C overexpressing cells suggesting that an increase in Lamin A overexpression marginally impacts TSA-induced nucleus stiffening. Thus, thickening of the nuclear lamina is insufficient to account for the full scale of nucleus stiffening after chromatin decompaction.

**Figure 4.**
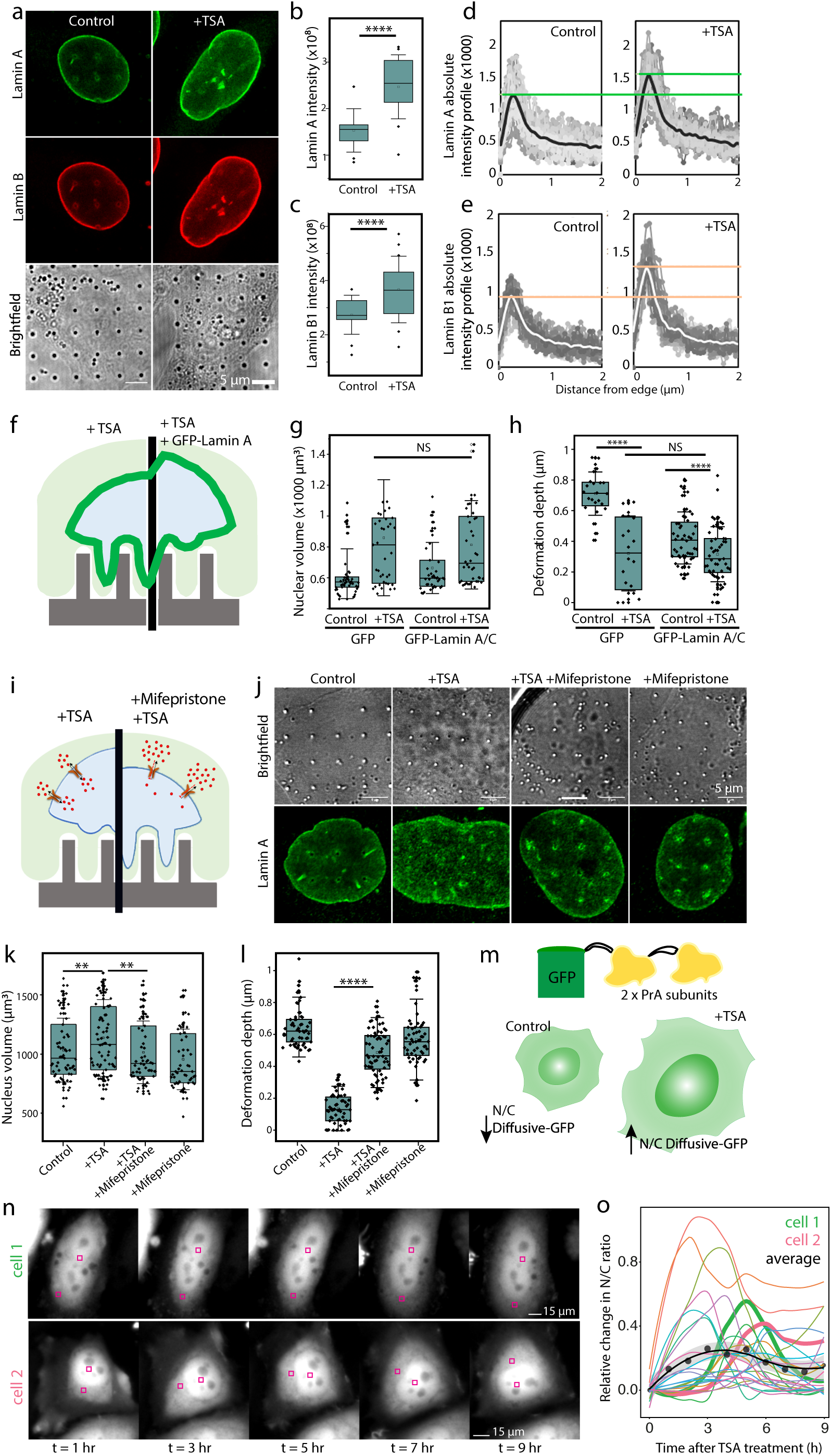
Nucleus stiffening after chromatin decompaction involves nuclear import and lamin recruitment. (a) Representative images of Lamin A and Lamin B in Control and TSA-treated cells. (b-c) Intensity of (b) Lamin A and (c) Lamin B1 measured from whole nuclei. (d-e) Intensity profile of (d) Lamin A and (e) Lamin B1 measured from the nuclear edge. (f) Schematic diagram of expected changes in nanopillar-induced deformation depth induced by Lamin A/C overexpression combined with acute TSA treatment. (g) Nucleus volume in response to overexpression of Lamin A/C and acute TSA treatment. (h) Nanopillar-induced deformation depth in response to overexpression of Lamin A/C and acute TSA treatment. (i) Schematic diagram of the effect of acute TSA and nuclear import inhibitor mifepristone treatment on nanopillar-induced deformation depth. (j) Representative images of cells on nanopillars after mifepristone and acute TSA treatment. (k) Nucleus volume in response to mifepristone and acute TSA treatment. (l) Nanopillar-induced deformation measured from nuclei after mifepristone and acute TSA treatment. (m) Schematic diagram showing the diffusive-GFP signal in the nucleus normalized to the cytoplasmic signal (N/C) in Control and TSA-treated nuclei. The diffusive-GFP fusion protein, consisting of the GFP protein and 2 subunits of the inert and purely diffusive bacterial protein Protein A (PrA). (n) Representative images of diffusive-GFP protein expressed by Hela cells and its localisation in response to TSA treatment time. (o) Relative change in N/C ratio of diffusive-GFP signal in response to TSA treatment. Each trace belongs to a single cell, with black dots and line denoting the average across all cells for each timepoint. Representative cells 1 and 2 are shown in green and pink, respectively. N = 25 cells analysed.

The other possible contribution to nucleus stiffness is the intranuclear pressure derived from the influx of molecules in response to chromatin decompaction, thereby providing a counterbalancing force exerted by the nuclear lamina. This hypothesis was supported by the correlation between changes in nucleus volume, nanopillar-induced deformation depth and Young’s modulus in response to TSA treatment time. To test this theory, we used mifepristone, an inhibitor of importin α/β-mediated transport^40^ (Fig. 4i). Compared to control, mifepristone alone did not change nucleus volume significantly but its combination with TSA treatment substantially reduced nucleus volume to the same level as the control (Fig. 4k). Deformation depth analysis confirmed that blocking of nuclear import using mifepristone dampened loss of nucleus deformability TSA treatment (Fig. 4l). We tested two other inhibitors of nuclear import, importazole and ivermectin and obtained similar results. Hela cells treated with both TSA and importazole or ivermectin for 4 hours enhanced H3K9ac intensity but did not influence nucleus volume or nanopillar-induced deformation depth compared to the control (Supplementary Figure 11). Additionally, we directly visualized enhancement of nuclear import in response to TSA treatment by tracking the localization of a 41 kDa diffusive protein construct fused to GFP (Fig. 4m). While cell-to-cell variability in relative intensity changes was high, individual cells exhibited a peak in the nuclear-to-cytoplasmic (N/C) ratio of the diffusive-GFP protein signal between 4-6 hours after TSA treatment (Fig. 4n-o, Supplementary Video 2). An average of an ensemble of cells showed a peak at 3 hours after TSA treatment, signifying an enhancement in nuclear import rate at similar timescale as nuclear stiffening (Fig. 4n-o). Interestingly, when we altered osmotic pressure by diluting cell culture media with distilled water, we observed an increase not only in nucleus volume and stiffness but also in H3K9ac density, indicating that material influx via nuclear import is induced during the decompaction of chromatin (Supplementary Figure 12).

Our results signify that immediately after receiving a signal for chromatin decompaction, there is influx of material (such as histone acetylation enzymes and molecules that may be required for chromatin decompaction or compensate the change in chromatin net charge) that leads to an acute increase in nucleus volume and turgidity. We have seen that this leads to increased nucleus stiffness; though what we are unable to decouple here is the contribution of the outward entropic pressure caused by chromatin decompaction and the release of chromatin condensation^41^. Based on previous work that has shown how large scale enzymatic decompaction of the chromatin causes rapid nuclear softening and swelling, we expected to observe that the increased nuclear volume would be a key response to accommodate chromatin expansion. However, to our knowledge our work is the first to study chromatin decompaction effects on nucleus mechanics in a completely intact cellular system. Nucleo-cytoplasmic feedback captured using our nanopillar platform thus has probably revealed a new mechanism of chromatin-based modulation of nucleus mechanics that relies on nuclear import. Not only that, the enhanced localization of lamin proteins to the nuclear envelope (presumably needed to protect the soft and now vulnerable genetic material) may restrict full nucleus expansion and dissipation of intranuclear pressures. This mechanoadaptation of the nucleus in response to chromatin decompaction seems to be relevant in other cellular processes too. T cell activation requires exit from quiescence and a new transcriptional program^42,43^ and is accompanied by a volumetric increase of the nucleus due to the rapid decompaction of chromatin needed for transcriptional accessibility^10^. Our data here implies that T cell activation leads to not merely stiffening of the whole T cell as previously reported^44^ but rigidification of the nucleus. This work thus implies that transient chromatin remodeling may have secondary consequences to important cell functions such as migration through regulation of nuclear stiffness.

### Correlation of nucleus deformability with chromatin decompaction is altered in malignant cancer cells

Mechanoadaptation of the nucleus should yield a pair of reciprocal phenomena: whilst acute chromatin decompaction causes nucleus stiffening (as shown in our work above), chromatin condensation should induce nuclear softening. Indeed previous reports show that elevation of global heterochromatin marked with H3K27me3 is required for the efficient migration of cells squeezing through transwells^9,45^ and between micropillar obstacles^46^, and that an increase in chromatin compaction can reshape and deform the nucleus effectively for confined migration^6,47^. We sought to test this phenomenon using breast cancer cell lines MCF7 and MDA-MB-231 (referred to as 231), which inherently differ in malignancy and metastatic potential, transwell migration rate, and epigenetic state^48^. Comparing against an untreated control, TSA treatment induced a significant increase in H3K9ac for all cell types (Fig. 5a) but a decrease in H3K27me3 level only in MCF7 cells (Fig. 5b). This is consistent with previous studies^49,50^ that report a decrease in heterochromatic H3K27me3 as a reciprocal effect to H3K9 hyperacetylation using TSA. The two types of histone modifications are linked by the EZH2 protein, the histone methyltransferase responsible for trimethylation of H3K27, binding to HDAC1^51^. Thus, suppression of HDAC1 activity by TSA not only increases histone acetylation (at both H3K9 and H3K27^52^) but also decreases trimethylation at H3K27. We have established in Hela cells that acute TSA treatment will swell the nucleus (Fig. 5c) and reduce transwell migration (Fig. 5d-e). MCF7 cells responded to TSA treatment in the same manner, albeit with no change in transwell migration due to the higher enlarged nucleus volume compared to Hela cells (Fig. 5d-e). Surprisingly, 231 cells exhibited no changes in either nucleus volume (Fig. 5c) or transwell migration (Fig. 5d) in response to TSA treatment, despite significant TSA-induced H3K9ac elevation against the control (Fig. 5a). We also observed enhancement in H3K27me3 levels after TSA treatment (Fig. 5b) contrasting with the TSA response in Hela and MCF7 cells. This is even more obvious in the ratio between H3K27me3 to H3K9ac intensity, where 231 cells exhibited extreme values for both untreated and TSA-treated nuclei compared to Hela or MCF7 (Supplementary Figure 13), indicating a bias in the highly malignant 231 cell type towards heterochromatin formation even under chromatin decompaction conditions. Concurrent with changes in nucleus volume and migration rate, TSA treatment significantly reduced deformation depth in MCF7 whilst inducing no change in 231 nucleus deformability (Fig. 5f-g). Young’s modulus measurements support the deformability measures (Supplementary Figure 14), indicating that 231 cells can maintain nucleus deformability even under chromatin decompaction challenges, likely through high heterochromatin formation and chromatin compaction.

**Figure 5.**
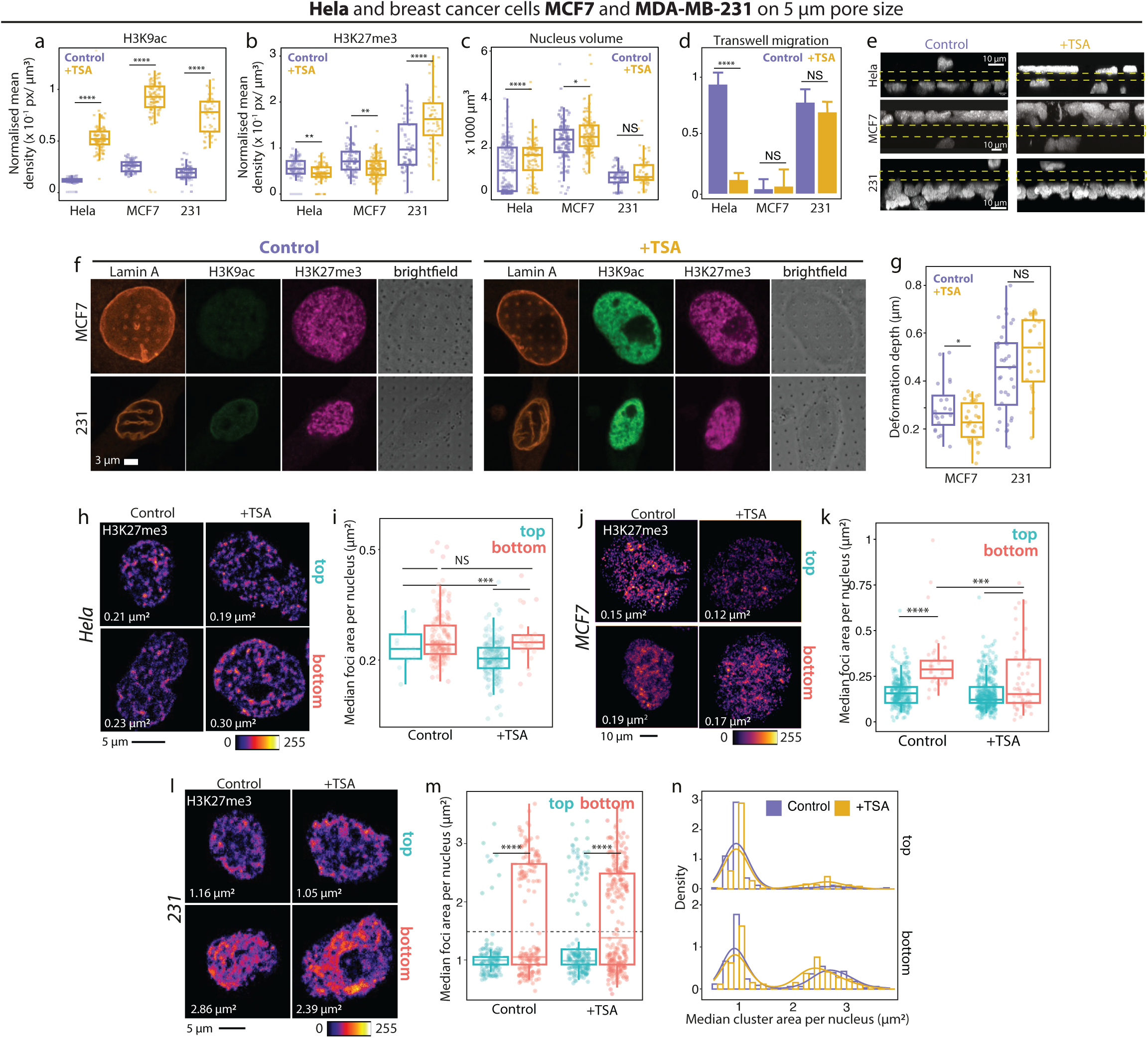
Malignant breast cancer cells exhibit differential nuclear deformability and chromatin compaction in response to acute chromatin decompaction. (a-c) Measures of (a) H3K9ac and (b) H3K27me3 density, and (c) nucleus volume in Hela, MCF7 and MDA-MB-231 (231) nuclei under control and acute TSA treatment. (d-e) Quantification and representative images of transwell migration of Hela and 231 cells under control and acute TSA-treated conditions. (f) Representative images of MCF7 and 231 stained against Lamin A, H3K9ac and H3K27me3 nuclei on nanopillars under control and acute TSA-treated conditions. (g) Nanopillar-induced deformation depth of MCF7 and 231 nuclei under control and acute TSA-treated conditions. (h-i) Representative images and quantification of size of H3K27me3 foci in Hela nuclei after acute TSA treatment. (j-k) Representative images and quantification of size of H3K27me3 foci in MCF7 nuclei after acute TSA treatment. (l-m) Representative images and quantification of size of H3K27me3 foci in 231 nuclei after acute TSA treatment. (n) Histogram shows the distribution of H3K27me3 foci in 231 nuclei according to size, with data binned at 0.2 μm. Images of H3K27me3 foci are enhanced for clear visualization of foci. Raw images of H3K27me3 are found in Supplementary Figure 15. Each dot represents one nucleus with the median of H3K27me3 foci size. Median area of H3K27me3 foci identified from the representative image is included.

We analyzed heterochromatin compaction in more detail by segmenting and measuring the size of H3K27me3 foci in nuclei on either side of a transwell (Supplementary Figure 15). We classified nuclei based on migration to the bottom of the transwell (‘bottom’, indicating a deformable/softer nucleus) versus those that remained on the top (‘top’, exhibiting a less deformable/stiffer nucleus). We and others^6^ postulate that larger H3K27me3 foci are more efficient at reducing chromatin physical occupancy to facilitate the deformability of the nucleus and migration of the cell. Regardless of TSA treatment, larger H3K27me3 foci were seen at the bottom compared to the top for Hela (Fig. 5h-i) and MCF7 (Fig. 5j-k). This reflects previous reports on the induction of global formation and local compaction of H3K27me3 by confined migration^9,46^. Super resolution images (using stimulated emission depletion) of Hela cells squeezing through a transwell show the same trend of larger and more distinct H3K27me3 foci when the nucleus segment is found at the bottom compared to the top (Supplementary Figure 16). Notably, H3K27me3 foci area in Hela and MCF7 cells decreased after TSA treatment within top cells – reflecting how chromatin decompaction can hinder heterochromatin foci compaction. In MCF7 cells at the bottom, we observed smaller foci size within bottom cells after TSA treatment cells which could indicate how Hela cells are more efficient than MCF7 cells at heterochromatin compaction even under TSA conditions. Our results imply that successful transwell migration requires the compaction of chromatin by increasing H3K27me3 foci formation, and that this process is obstructed by TSA-induced chromatin decompaction.

In 231 cells, H3K27me3 were compacted into extremely large foci with area that was an order of magnitude larger than those found in MCF7 or Hela cells (Fig. 5l-m). A detailed look at the distribution of the H3K27me3 foci size showed two clearly segregated subpopulations, where one subpopulation was observed for foci with area larger than 1.5 µm^2^ (‘large foci’, Fig. 5n). For cells that remained at the top, TSA showed no discernible effect on the large foci size. However, for cells that migrated to the bottom, the large foci significantly shrank after TSA treatment (p-value = 2.8 x 10-4) though this appears to have no influence on nucleus deformability. The lack of change in chromatin compaction correlated with maintenance of deformability in response to TSA treatment, strongly indicating to us the efficient reshaping of the nucleus by 231 cells via heterochromatin compaction.

We then examined pairs of ovarian and lung cell lines with different malignancy levels. Both pairs of ovarian and lung cancer cell lines responded to TSA treatment by increasing H3K9ac levels (Supplementary Figure 17). Mirroring the response of 231, TSA treatment only induced an elevation in H3K27me3, which were highly correlated to the maintenance of nucleus volume and transwell migration in highly malignant HEYA8 (ovarian) and HT1080 (lung) cells compared to poorly malignant PEO1 (ovarian) and A549 (lung) cells, respectively. Nanopillar-induced deformation depth exhibited similar results to what we observed in breast cancer cell lines, where TSA treatment decreased the deformability of PEO1 and A549 but not HEYA8 and HT1080. In transwell assays, heterochromatin H3K27me3 in highly malignant cells reduced in size (for ovarian cancer cells) and intensity (for lung cancer cells) but at a smaller degree than observed in poorly malignant cells. Essentially, the dichotomy in acute response to TSA between poorly and highly malignant cells were also seen in ovarian and lung cancer cell lines.

Chromatin organization and modification are overtly changed during cancer progression, with the elevation of H3K27 methyltransferase EZH2 linked to a transcriptional program that promotes cell invasiveness and anchorage independent growth in aggressive prostate and breast cancers^53,54^. However, the non-transcriptional, mechanical role of chromatin in pro-malignancy changes is underappreciated. Analysis of poorly malignant breast, ovarian and lung cancer cells consistently exhibited that acute decompaction of the chromatin effectively opposes heterochromatin foci formation and permits nucleus swelling that hinders cell migration. In contrast, the capacity of highly malignant cells to counter chromatin decompaction through heterochromatin formation presumably underlying nuclear reshaping and efficient migration under confinement (Fig. 7). This represents a mechanism in which malignant cancer cells can be mechanically favored over poorly malignant ones, by providing a swifter means of achieving characteristics promoting a malignant phenotype compared to the process of genomic instability^55^. Whether the phenomenon we observed here is a cause or consequence of cancer-associated mutations is an interesting connection to pursue further and could open new targets for anti-metastatic drug discovery. Indeed, RNA-seq has recently revealed that histone-modifying genes (including those encoding for histone deacetylases and histone methyltransferases) are frequently altered and may drive the development of non-Hodgkin lymphoma^56^. Our work underlines how chromatin organization is an important dictator of transient nuclear mechanical properties that impinge on cell behaviors in healthy and diseased states.

**Figure 6.**
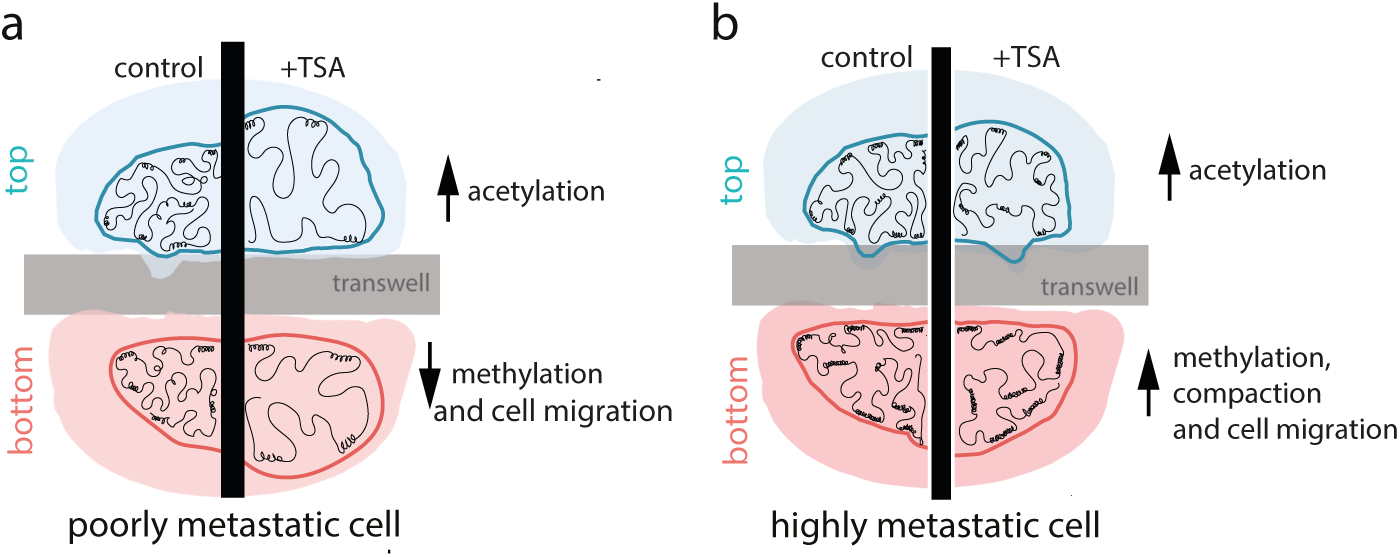
Schematic figure showing response of poorly and highly malignant cells to acute chromatin decompaction.

## MATERIALS AND METHODS

### Nanopillar fabrication

Nanopillars (500 nm diameter, 1.5 µm height, 3-5 µm pitch) used in the work were fabricated using techniques previously described. Briefly, quartz chip was spin-coated with polymethylmethacrylate (300 nm thickness) and conductive coating (AR-PC 5090.02, Allresist). Nanopatterns were written using electron beam lithography (FEI Helios Nanolab 650), with exposed areas removed using isopropanol:methylisobutylketone (3:1) solution. Afterwards, a Cr mask (80 nm thickness) was deposited on the surface using thermal evaporator (UNIVEX 250 Benchtop) with the leftover PMMA layer lifted off with acetone. To reveal nanopillars, reactive ion etching using CF_4_ and CHF_3_ (Oxford Plasmalab 80) was performed. SEM was used to determine quality and quantify dimensions of resulting nanopillars. Prior to cell seeding, samples were cleaned with Cr etchant (Transene) and O2 plasma.

### Cell culture, biochemical treatments, immunostaining, overexpression and knockdown

Hela, MCF7 and MDAMD231 cells were obtained from ATCC. PEO1 and HEYA8 ovarian cancer cell lines were kind gifts from Prof. Joanne Ngeow. A549 and HT1080 lung cancer cell lines were kind gifts from the Mechanobiology Institute. Hela, A549 and HT1080 cells were maintained in media comprising DMEM (high glucose, with L-glutamine and sodium pyruvate) with 10% fetal bovine serum (FBS, Gibco) and 100 U ml^−1^ penicillin and 100 mg ml^−1^ streptomycin (Sigma-Aldrich). MDA-MB-231 and MCF7 cells were maintained in media comprising DMEM (high glucose, with L-glutamine) with 10% FBS and 100 U ml^−1^ penicillin and 100 mg ml^−1^ streptomycin. PEO1 and HEYA8 cells were grown in RPMI supplemented with 10% FBS and 100 U ml^−1^ penicillin and 100 mg ml^−1^ streptomycin. All cells were trypsinized using 0.05% trypsin in EDTA (Gibco). Cells were grown and maintained in standard cell culture conditions of 37°C and 5% CO_2_.

Nanopillars or flat coverslips were first UV-sterilized before coating with human fibronectin (∼0.1 mg/cm^2^, Sigma-Aldrich) before seeding with cells. To induce chromatin decompaction, cells were treated with TSA (Sigma-Aldrich) or valproic acid (Sigma-Aldrich) at indicated concentrations and timescales. Cells were also treated with AA (30 μM, Sigma-Aldrich) to inhibit histone acetylation or mifepristone (30 μM, Sigma-Aldrich) to inhibit nuclear import.

For AKTIP knockdown, HeLa cells at 70-80% confluency were transfected with 100 pM of siRNAs (siControl or siAKTIP) using lipofectamine 3000 (Invitrogen). Cells were grown for 72 hours post-transfection to ensure efficient gene knockdown before cells were used for further experiments. For overexpression of lamins, HeLa cells were transfected with GFP-LaminA/C construct or GFP alone using lipofectamine 3000, followed by 16 hours culture before cells were trypsinised and re-plated on nanopillars.

### Transwell migration assay

Cells on a 6-well plate were left untreated or pre-treated with TSA, plated on top of a Transwell polyester membrane (24 well plate size, Corning) then allowed to migrate for a specified amount of time before fixation. For Hela, cells were pre-treated for 90 mins then allowed to migrate for 90 mins. For breast cancer cells, all cells were pre-treated for 1 hour then allowed to migrate for 3 hours, for a total treatment time of 4 hours. For ovarian and lung cancer cells, cells were seeded on transwells and allowed to migrate for 4 hours followed by TSA treatment and an additional 4 hours of migration time. For Hela, MDA-MB-231, A549 and HT1080 cells, 5 µm pore size transwell was used, while a pore size of 8 µm was used for MCF7, PEO1 and HEYA8 cells.

### Immunostaining and confocal imaging

Cells on nanopillars, coverslips or transwells were fixed with 4% paraformaldehyde for 20 mins at room temperature. Samples were then permeabilized with 0.5% Triton-X (Sigma) in phosphate buffered saline (PBS) for 6 mins. After washing thrice with PBS, cells were blocked with 5% BSA (blocking solution) for 1 hour at room temperature. This was followed by incubation for 1.5 hours at room temperature with the primary antibody of choice: anti-LaminA (ab26300, Abcam), anti-LaminA/C (4777, Cell Signaling Tech), anti-H3K9ac (ab10812, Abcam and GTX630554, GeneTex), Anti-LaminB1 (ab16048, abcam), anti-AKTIP (WH0064400M2, clone 2A11, Sigma Aldrich), anti-H3K27me3 (GTX121184, GeneTex). Cells were washed with PBS then incubated with the corresponding Alexa Fluor-conjugated (488, 546 or 647 fluorophores) secondary antibody (1:500; Life Technologies). The nuclei were labeled with DAPI. For transwell assays with tricolor stains, anti-Lamin A antibody (ab26300, Abcam) was conjugated with a 550 fluorophore (CoraLite conjugation kit, Proteintech) then placed on cells for 1 hour at room temperature. Imaging of cells on nanopillars was performed using Zeiss LSM 800 confocal microscope using a 100X objective (NA = 1.4, oil immersion) and Z-step of 0.15-0.5 μm. Images of nuclei adhered on flat coverslips and transwells were captured using a 63X objective (NA = 1.2, oil immersion) and a Z-step of 0.4um. Imaging conditions were kept similar across all samples within individual experiments. Image analysis is detailed in the supplementary material.

### Atomic Force Microscopy

AFM indentation experiments were performed using a Nanowizard 4 BioScience AFM (JPK Instruments, Germany) at 37°C and in a 5% CO2 environment. The probe consisted of a 4.5 µm diameter polystyrene bead attached to a silicon nitride cantilever (Novascan Technologies, Inc., Ames, IA, USA). The spring constant of the cantilever used was determined by the thermal tune method in liquid and in the range of 0.03 ± 0.003 N/m. Indentations were performed at the nuclear regions of randomly selected cells at 5 μm/s probe velocity and 1 nN trigger force within a 2 x 2 μm^2^ area and 5 x 5 pixel resolution. An average of 30 cells were tested for each condition and at least 750 force curves were obtained for analysis to estimate the Young’s modulus. These values were calculated for each recorded curve using JPK Data Processing Software (JPK Instruments, Germany), which employs a Hertz’s contact model for spherical indenters (diameter of 4.5 µm; Poisson’s ratio of 0.5) fitted to the approach curves. Representative force-depth response curves and pointwise modulus plots are shown in Supplementary Figures 18 and 19.

### Timelapse using Lamin A chromobody

The Lamin A-TagGFP2 chromobody plasmid (ChromoTek), which produces TagGFP2-tagged intracellular functional antibody against Lamin A, was used to track nuclear deformation of Hela cells using live cell microscopy. Hela cells seeded on nanopillars were transfected with the plasmid using lipofectamine 2000 (LifeTechnologies). After 36-48 hours, live imaging was performed using 100X objective (NA = 1.45, oil immersion) on a spinning disc confocal microscope built around a Nikon Ti2 inverted microscope. Hela cells were treated with TSA (250 ng/μL) then tracked every hour for 4 hours.

### Timelapse imaging using Diffusive-GFP fusion protein

Hela cells grown on glass bottom dish were transfected with the plasmid expressing the 41 kDa Diffusive-GFP construct (addgene # 201335) using Lipofectamine 2000 (ThermoFisher). The diffusive-GFP fusion protein consists of the GFP protein linked to 2 repeats of the inert and diffusive bacterial protein A (7 kDa)^57^. Twenty four hours after transfection, cells were treated with TSA and immediately imaged using a Nikon Eclipse Ti-2 epifluorescence microscope with a 40x objective (numerical aperture = 0.6) and LED illumination. Images were acquired every hour. For each nucleus, the mean intensity of GFP was subtracted from the background, which was obtained from an area without cells. The mean intensity of GFP in the cell was also subtracted from the mean intensity in the background and was subsequently used to obtain the nucleus-cytoplasmic (N/C) ratio of diffusive GFP signal, as previously described^57^. To normalize cell-to-cell variability data, we calculate the relative change in N/C GFP ratio at each timepoint relative to the first timepoint.

### H3K27me3 foci analysis

H3K27me3 foci analysis follows the methodology provided by McQuin et al. for CellProfiler^58^. Images of nuclei stained against H3K27me3 across different Z planes were obtained and projected at the midpoint of the nucleus (using maximum projection method). Projected images of H3K27me3 were then enhanced to increase signal to noise ratio. Thereafter, enhanced images were thresholded using an adaptive Otsu strategy with three-class thresholding to segment H3K27me3 foci. Segmented foci were then used to calculate H3K27me3 foci size. High resolution microscopy of H3K27me3 foci were performed on Hela cells by imaging on the Abberior STEDycon microscope (Abberior Instruments GmbH) equipped with pulsed STED line at 775 nm, excitation lasers at 485 and 640nm with 100 nm pixel size and 100x oil immersion objective (NA = 1.4).

### Statistical analysis

Data in box plots show the interquartile range, while bar charts show mean ± s.e.m. unless otherwise stated in the figure legends. Data analyzed were collected from at least three biological replicates N ≥ 3. One-way ANOVA with Tukey’s post-hoc test was used for significance testing for all Figures except Figure 5, which used Two-way ANOVA with Tukey’s post-hoc test. p < 0.05 was considered significant. Analyses were performed using OriginPro (v9.0) and R (v4.1.1) through its graphical interface RStudio (v2021.09.0). Line graphs and bar plots were generated using Origin and ggplot2 package (v3.4.0) for R. NS indicates no statistical significance).

## ACKNOWLEDGEMENTS

This research was supported by funding from the Singapore Ministry of Education (RG145/18, RG93/22, RT3/22 to WZ; MOE-MOET32020-0001 to WZ and AL), Ministry of Healthy (Healthy Longevity Catalyst Awards (MOH-001192-01 to WZ)), NTUitive (NGF-2021-10-026 to WZ), Nanyang Technological University (Start-Up Grant to WZ), the Institute for Digital Molecular Analytics and Science (to WZ), Mechanobiology Institute Seed Grant (to CTL), AIRC (IG-24614) to IS and NBFC support funded by the National Recovery and Resilience Plan (NRRP), Mission 4 Component 2 Investment 1.4 - Call for tender No. 3138 of 16 December 2021, rectified by Decree n.3175 of 18 December 2021 of Italian Ministry of University and Research funded by the European Union - NextGenerationEU; Award Number: Pro, Project code CN_00000033, Concession Decree No. 1034 of 17 June 2022 adopted by the Italian Ministry of University and Research, Project title “National Biodiversity Future Center - NBFC” Sapienza CN5-Spoke 7 (to IS). IS is affiliated to the following institutions: School of Biological Sciences, Nanyang Technological University (NTU) Singapore, NTU Institute of Structural Biology (NISB) Singapore, CNR Institute of Molecular Biology and Pathology, Rome, Italy, and Instituto Pasteur Fondazione Cenci Bolognetti, Rome, Italy. MFAC acknowledges the support of the NTU Presidential Postdoctoral Fellowship. We thank Prof. Joanne Ngeow and Siao Ting Chong for fruitful discussions on breast cancer cell responses. We thank Zhiming Koh for his assistance with cell culture. The authors also acknowledge the support of the School of Biological Sciences and the School of Chemistry, Chemical Engineering and Biotechnology in NTU for the confocal microscope access, and the Nanyang NanoFabrication Center (N2FC), the Facility for Analysis, Characterization, Testing and Simulation (FACTS), and the Centre of Disruptive Photonic Technologies (CDPT) in NTU for supporting the nanopillar chip fabrication and characterization, as well as the NISB Cryo-EM lab in NTU where electron microscopy data collection was performed.

## AUTHOR CONTRIBUTIONS

Conceptualization, AM, GVS, IS and WZ. Investigation, AM, MFAC, RB, YZ, NQ, MK, BV, NMH and BH. Formal analysis, AM, MFAC. Writing – Original Draft, MFAC; Writing – Visualization, AM, MFAC; Writing – Review & Editing, AM, MFAC, GVS, IS, WZ. Project administration and funding acquisition, AL, CTL, GVS, IS and WZ.

