## Supplementary Figures for "Acute chromatin decompaction stiffens the nucleus as revealed by nanopillar-induced nuclear deformation in cells"

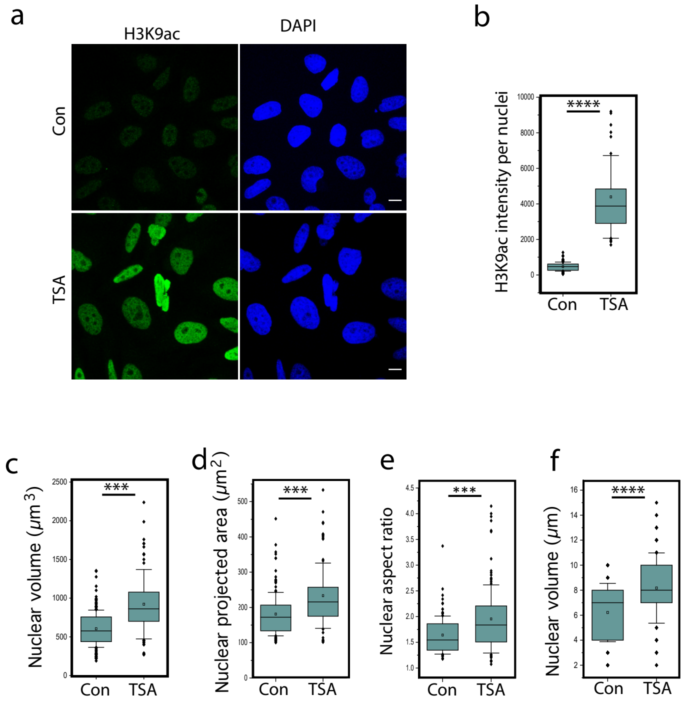


Supplementary Figure 1. Chromatin decompaction induced by trichostatin A (TSA) leads to changes in nuclear morphology. Hela cells were grown on flat coverslips and treated with TSA for 4 hours. (a) Representative images of Hela cells stained against H3K9ac and DAPI after treatment. Scale bar = 10 $\mu$m. (b) Box plot showing H3K9ac levels in Hela nuclei. Box plots showing geometrical properties of the nuclei such as (c) nuclear volume (d) nuclear spread area, (e) nuclear aspect ratio and (f) nuclear height. Con denotes untreated control sample. Cells were treated with TSA for 4 hours.

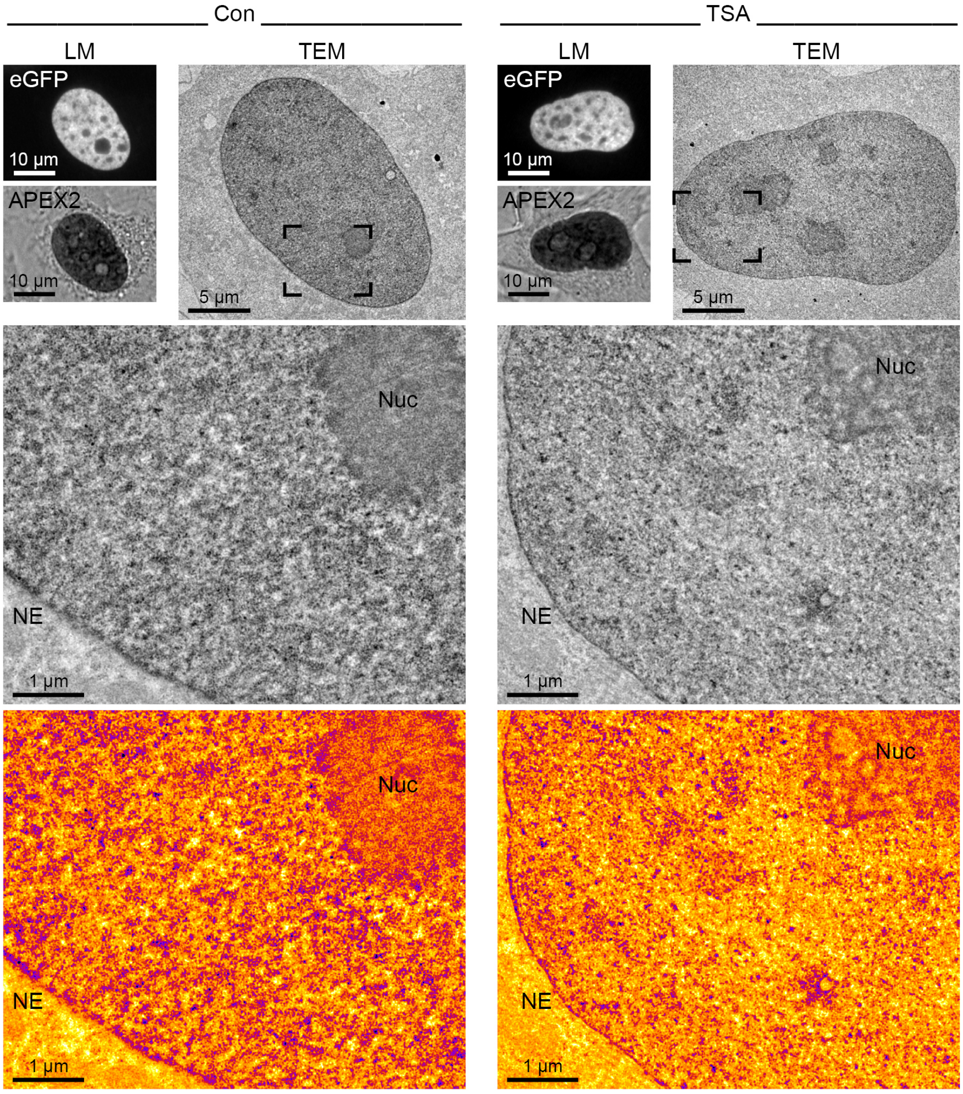


Supplementary Figure 2. Effect of TSA on chromatin structure. Transmission electron microscopy (TEM) shows chromatin decompaction after TSA treatment (TSA, 3.5 h) along the nuclear envelope (NE), around nucleoli (Nuc), and in most other regions of the nucleus compared to control cells (Con). In the heat map images (bottom row) purple represents most compact chromatin, yellow represents most decondensed chromatin. Cells with similar expression levels of eGFP-APEX2-H2B were chosen as confirmed by light microscopy (LM).

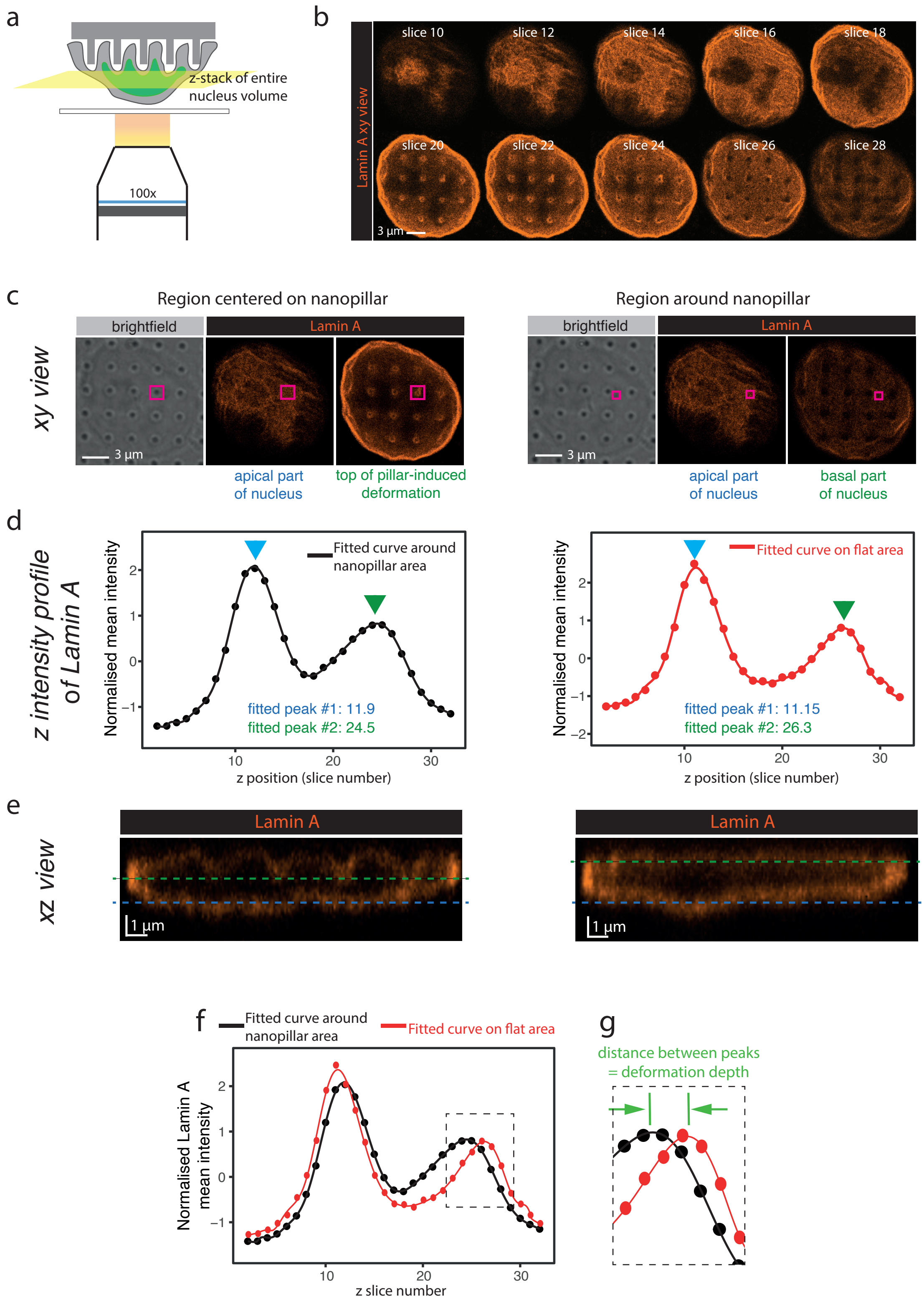


Supplementary Figure 3. Measurement of deformation depth induced by nanopillars using changes in Lamin A intensity. (a) Schematic of high resolution confocal imaging performed on cells on nanopillar substrates that were inverted onto coverslips. (b) Montage of xy view of Lamin A images across the entire nucleus. Slices 20 and 22 show clear deformation of nanopillars (seen as rings) on the Lamin A. (c) Representative images of nuclei with regions of interest centered on nanopillars or drawn around nanopillars. (d) Raw and fitted intensity profile of Lamin A along the Z direction. Blue arrows denote the first peak that identifies the apical part of the nucleus. Green arrows denote the second peak that identifies either the top of the nanopillar-induced deformation of the nucleus, or the basal aspect of the nucleus. In the region centered around the nanopillar, the second maximum is closer to the first maximum because of how the nanopillars indent the nuclear envelope from the bottom towards the top. The deformation depth is calculated by obtaining the difference in the second peak between the nanopillar area and surrounding area. (e) Cross section or xz view of the nucleus as viewed when centered on nanopillars or around nanopillars, with blue and green dashes representing the two peaks of the Z intensity profile. (f) Overlay of intensity profile of Lamin A along Z direction for nanopillar area and flat area. (g) Inset shows that the distance between the two peaks of Lamin A intensity profile is the calculated deformation depth.

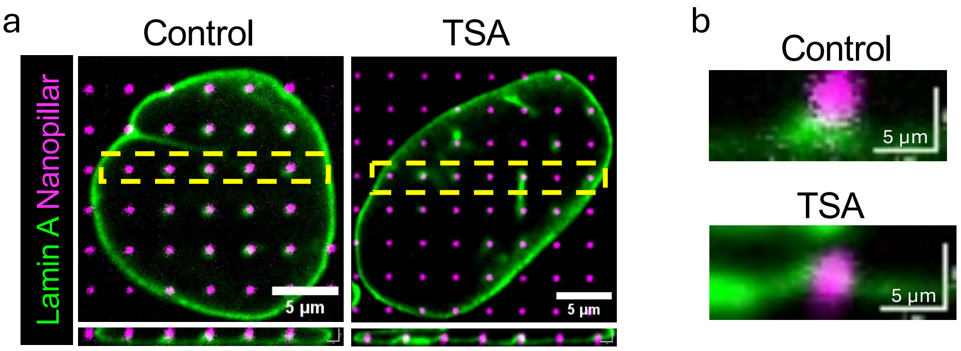


Supplementary Figure 4. Images of cells and nanopillars prior to expansion microscopy. (a) Shows overview of cell in xy view and an array of pillars in xz view (denoted by yellow box). (b) Representative xz view from the center of a nanopillar. The same cells are shown in Figure 1 after expansion microscopy.

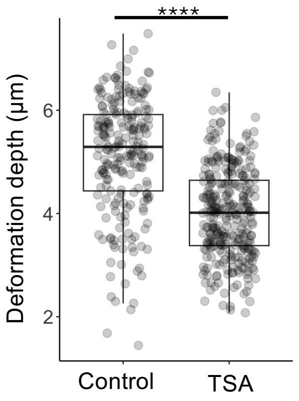


Supplementary Figure 5. Deformation depth measured from individual nanopillar xz views acquired via expansion microscopy.

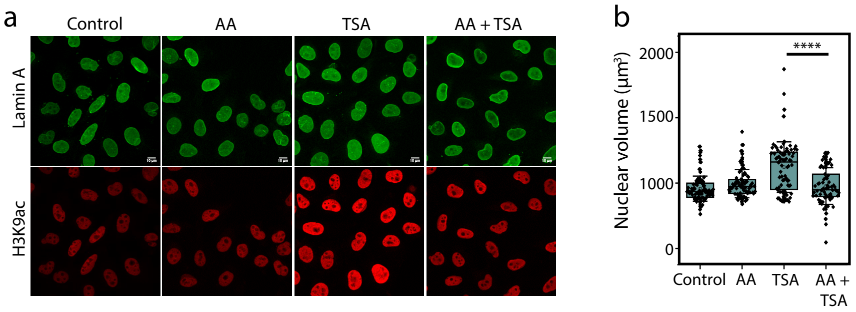


Supplementary Figure 6. (a) Representative image of Hela cells cultured on coverslips in untreated condition (control), or treated with TSA, anacardic acid (histone acetyltransferase inhibitor) or both. Nuclei were staineds against Lamin A (green) and H3K9ac (red). (b) Boxplots showing the nuclear volumes after different treatments.

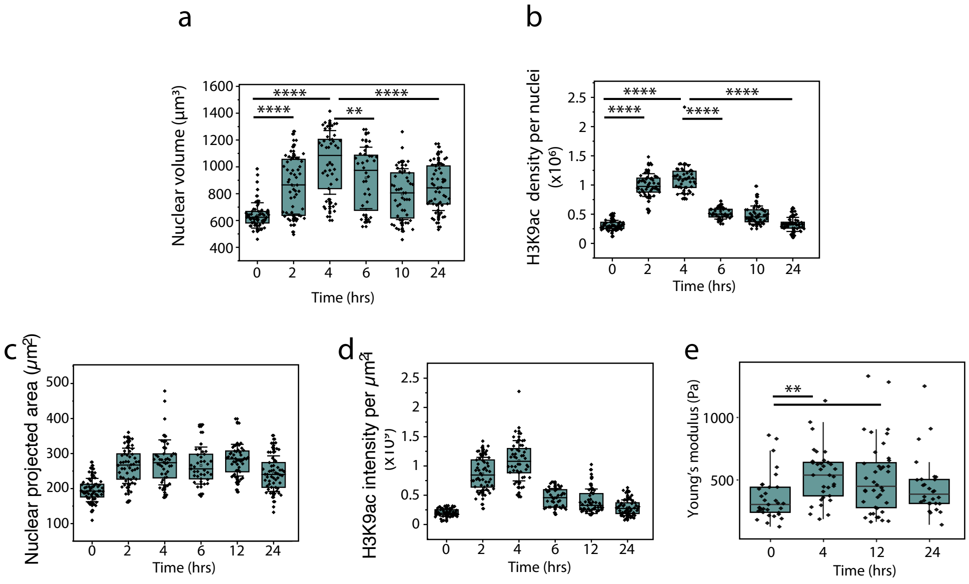


Supplementary Figure 7. Chromatin decompaction-mediated stiffening of the nucleus induced by TSA treatment is time-dependent. Box plots showing the (a) nucleus volume, (b) nuclear H3K9ac density, (c) nucleus area, and (d) nuclear intensity and (e) Young's modulus of the nucleus in Hela cells treated with TSA at different durations.

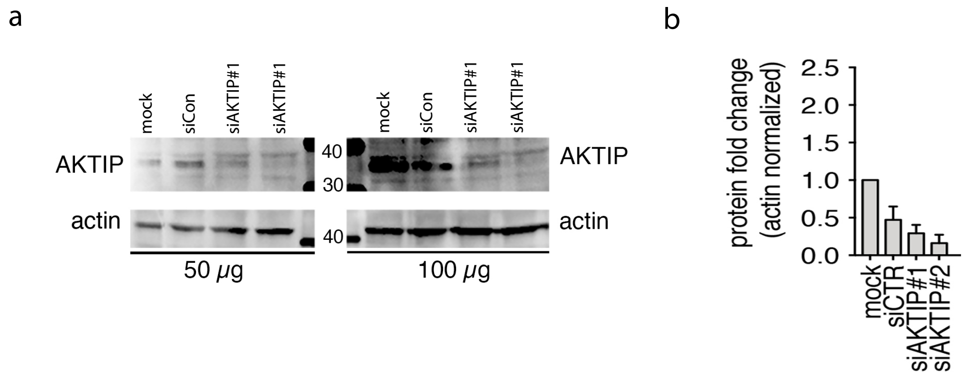


Supplementary Figure 8. siRNA-mediated knockdown of AKTIP induces increase in nuclear size and stiffness. (a) Western blots showing the knockdown of AKTIP protein using siRNA against AKTIP. Lysates of Hela cells mock transfected (mock) or transfected with scramble siRNA (siControl) or siRNAs against AKTIP (siAKTIP) were immunoblotted for AKTIP and actin. (b) Quantications of AKTIP protein level nomalised to actin.

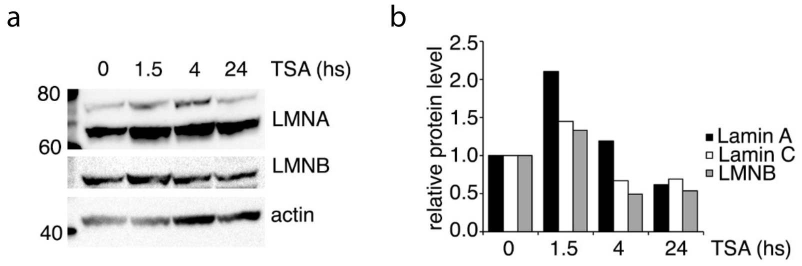


Supplementary Figure 9. HDAC inhibition by TSA induces elevated Lamin A levels at short timescales. (a) Western blot and (b) its densitometric quantification showing the expression levels of Lamin A (upper band in Western blot), Lamin C (lower band in Western blot) and Lamin B1 after different duration of TSA treatment.

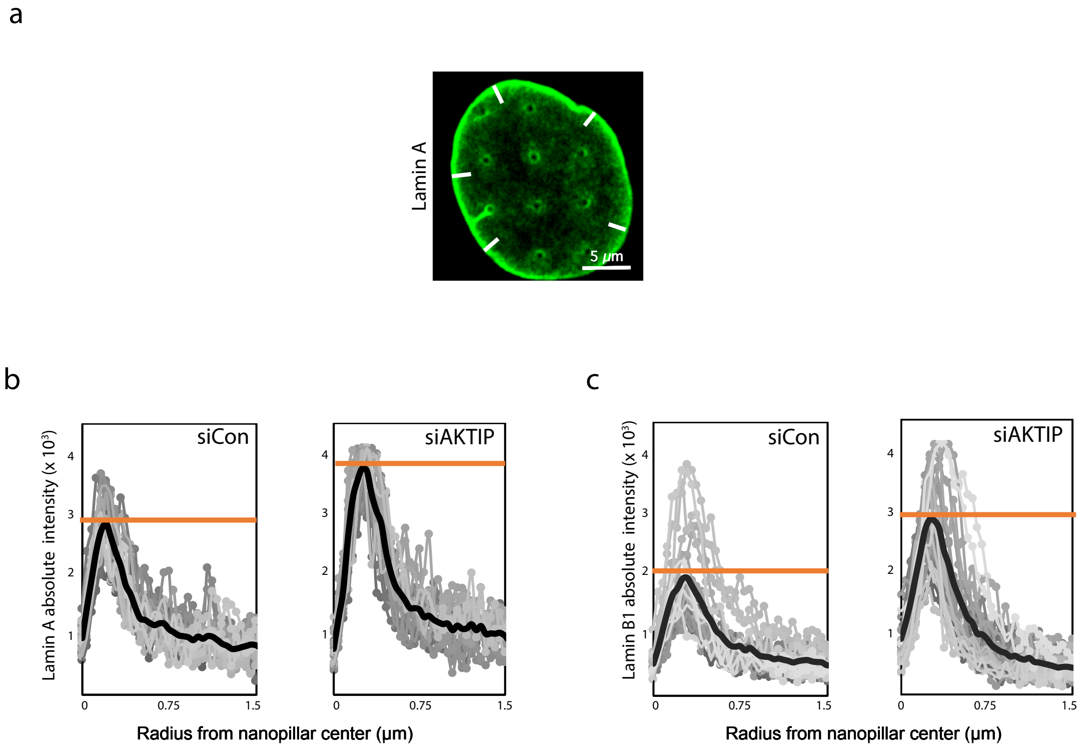


Supplementary Figure 10. AKTIP-depletion mediated nuclear stiffening involves recruitment of Lamins at the nuclear lamina. (a) Representative image showing the linear ROIs (white bar) at the periphery or edge of the nucleus, as denoted by the increased lamina intensity. ROI length used to measure the linear intensity profile of lamins is 1.5 $\mu$m. Scale bar = 5 $\mu$m. The linear intensity profiles of (b) Lamin A and (c) Lamin B1 at the nuclear peripheral lamina in control and AKTIP depleted cells.

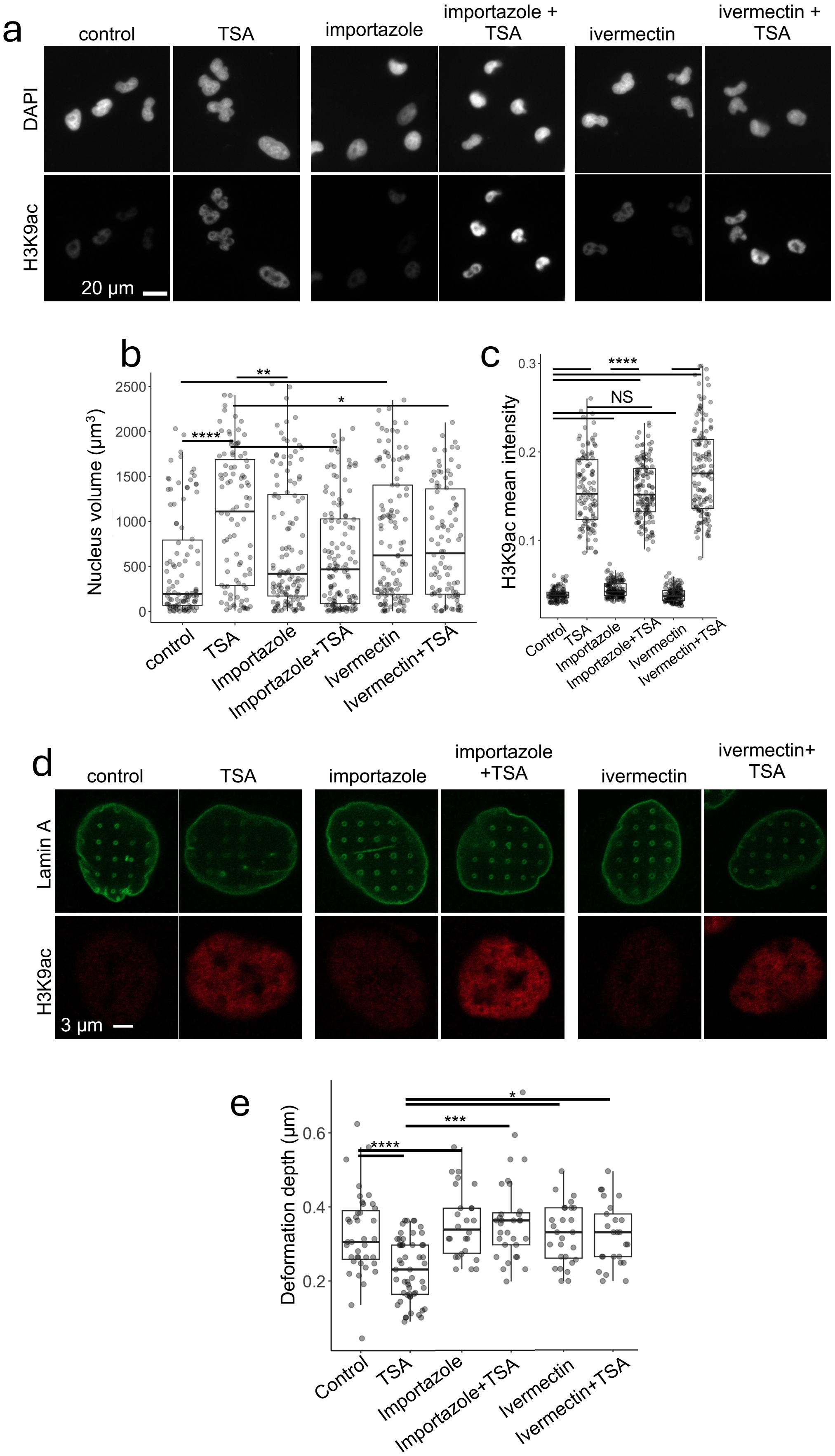


Supplementary Figure 11. Nuclear import inhibition using importazole and ivermectin recovers the deformability of the nucleus after TSA treatment. (a) Representative images, (b) nucleus volume measurements and (c) H3K9ac intensity of Hela cells grown on flat coverslips and treated with TSA and nuclear import inhibitors. (d) Representative images and (e) deformation depth of Hela cells grown on nanopillars and treated with TSA and nuclear import inhibitors.

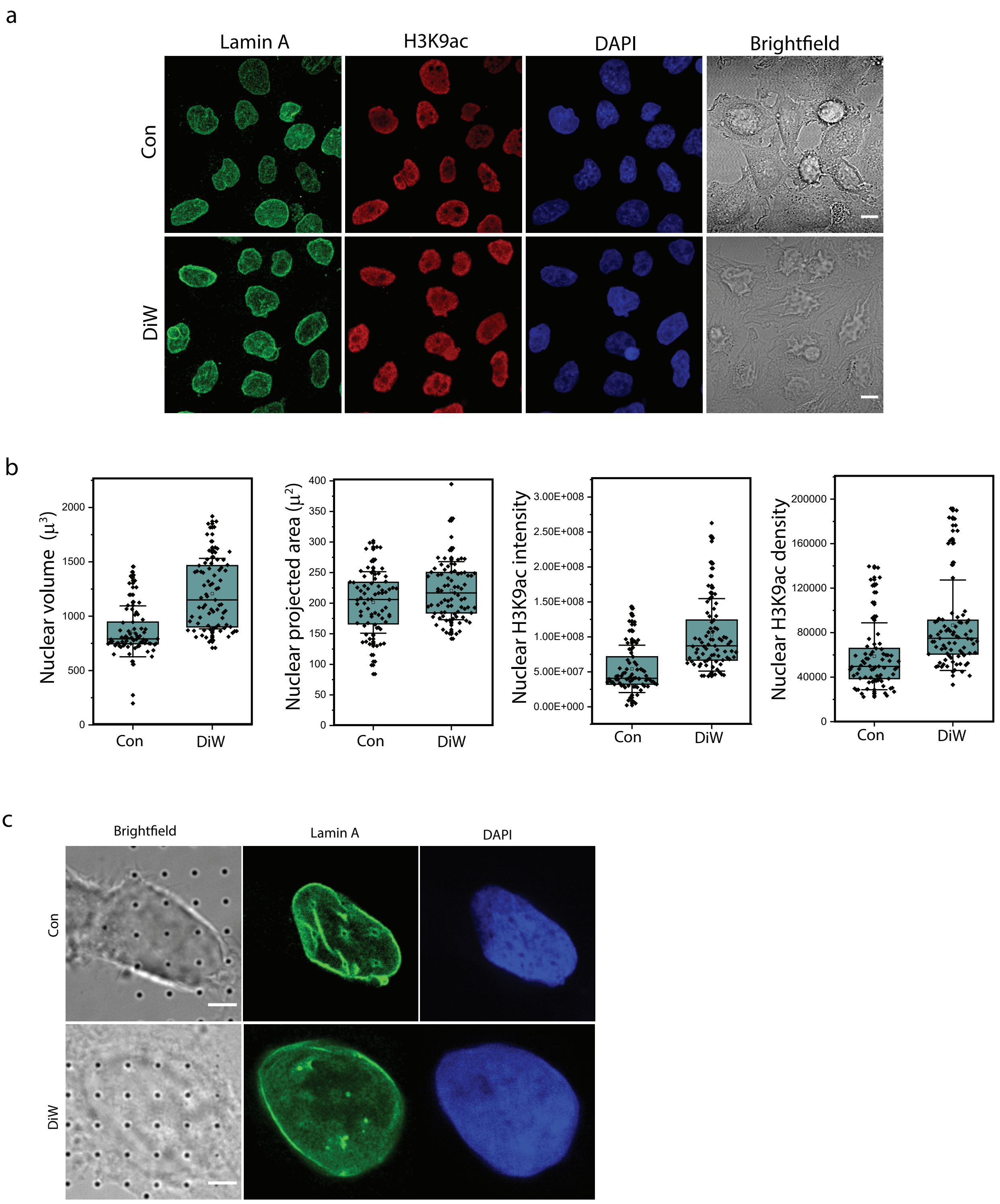


Supplementary Figure 12. Nucleus properties of Hela cells after treatment with distilled water (DiW). (a) Representative images of Hela cells with or without treatment of DiW for 3 mins. Nuclei were stained for Lamin A (green), H3K9ac (red), DAPI (blue). Scale bar = 30 $\mu$m. (b) Box plots showing the nuclear volume, area, H3K9ac intensity and H3K9ac density after DiW treatment. (c) Representative images of Hela cells on nanopillars after treatment with DiW. Nuclei treated with DiW show poor pattern formation against nanopillars. Scale bar = 5 $\mu$m.

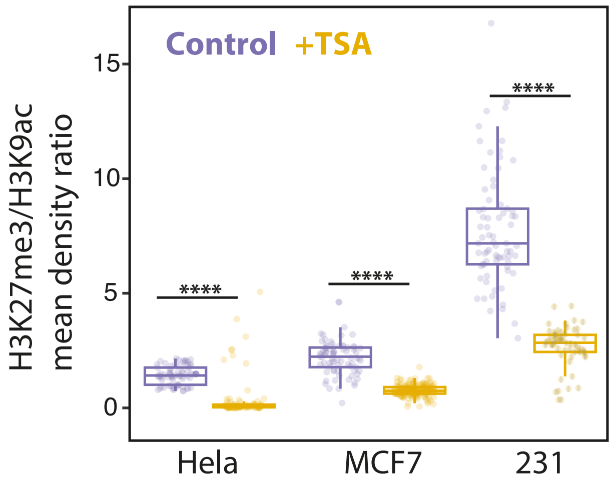


Supplementary Figure 13. Ratio of H3K27me3 to H3K9ac intensity in Hela, MCF7 and 231 cells.

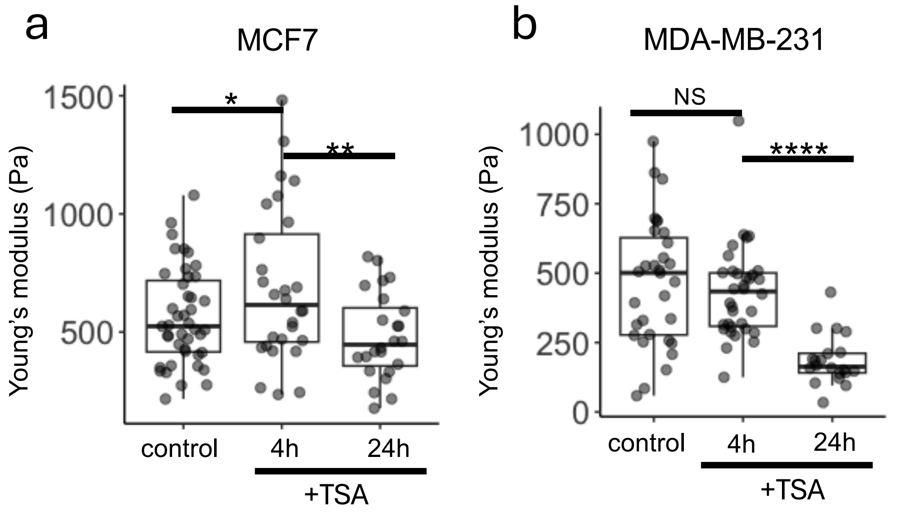


Supplementary Figure 14. AFM measurements of the nuclei of (a) MCF7 and (b) MDA-MB-231 in response to TSA treatment over time. Young's moduli were obtained by fitting to a fixed indentation depth of 600 nm.

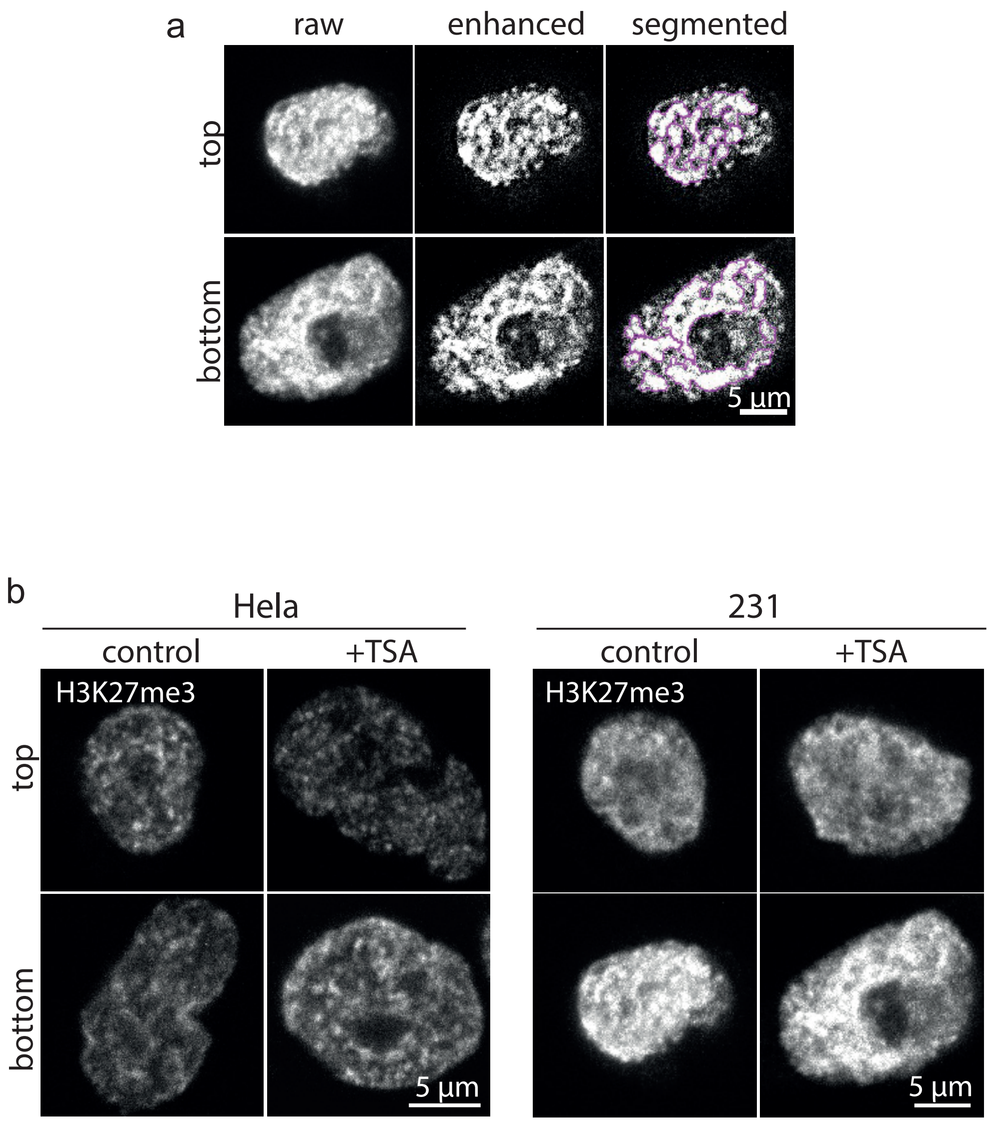


Supplementary Figure 15. Identification of H3K27me3 foci. (a) Images of H3K27me3 were first enhanced to subtract unwanted background from cells. Enhanced images were then used to segment foci using an adaptive thresholding strategy. (b) Raw images of H3K27me3 from Hela and 231 cells. These images correspond to enhanced images shown in Figure 5.

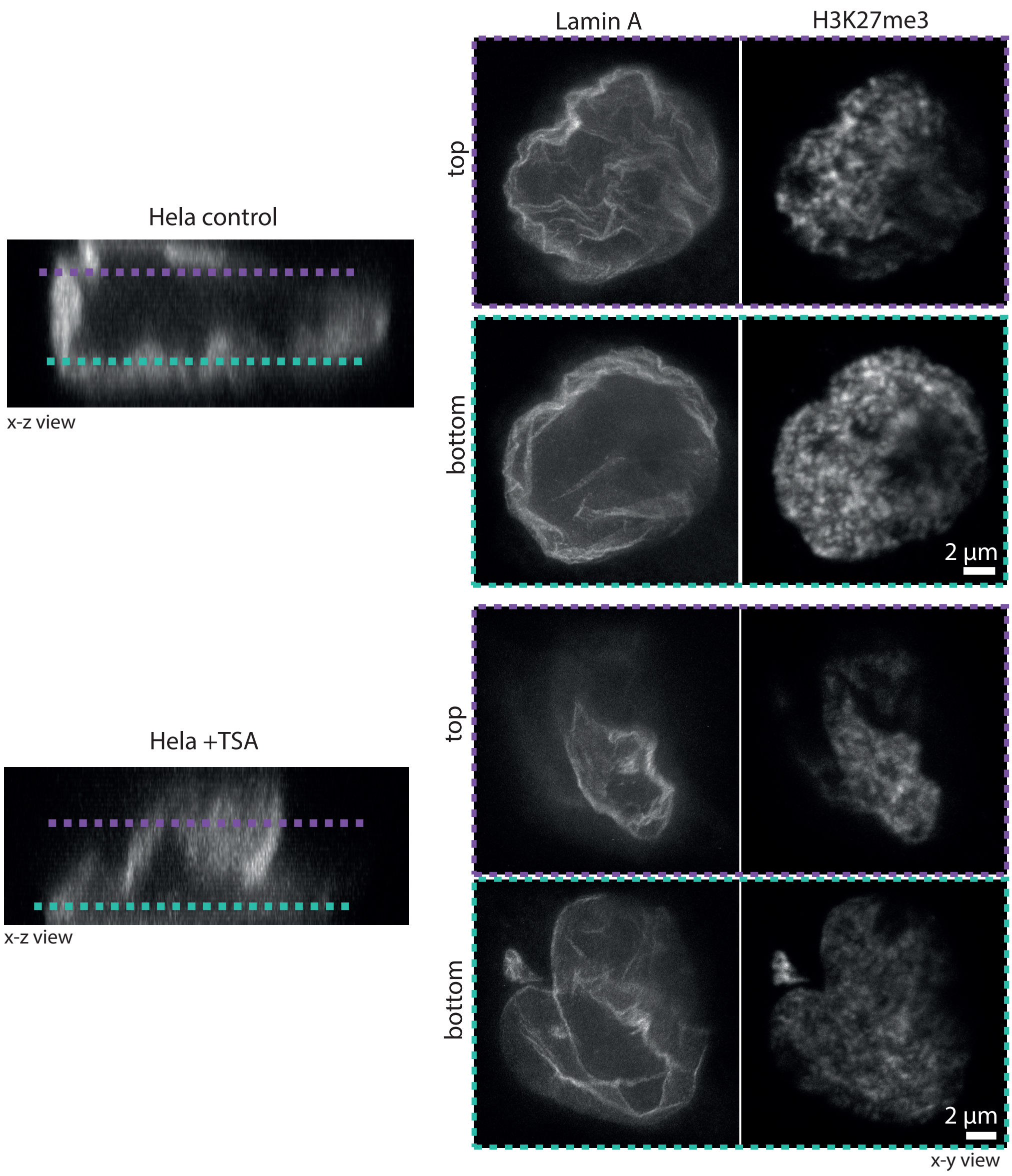


Supplementary Figure 16. Fluorescence nanoscopy of single Hela nuclei squeezing through a transwell pore. Stimulated emission depletion microscopy (STED) was used to image untreated and TSA-treated Hela cells stained against Lamin A and H3K27me3.

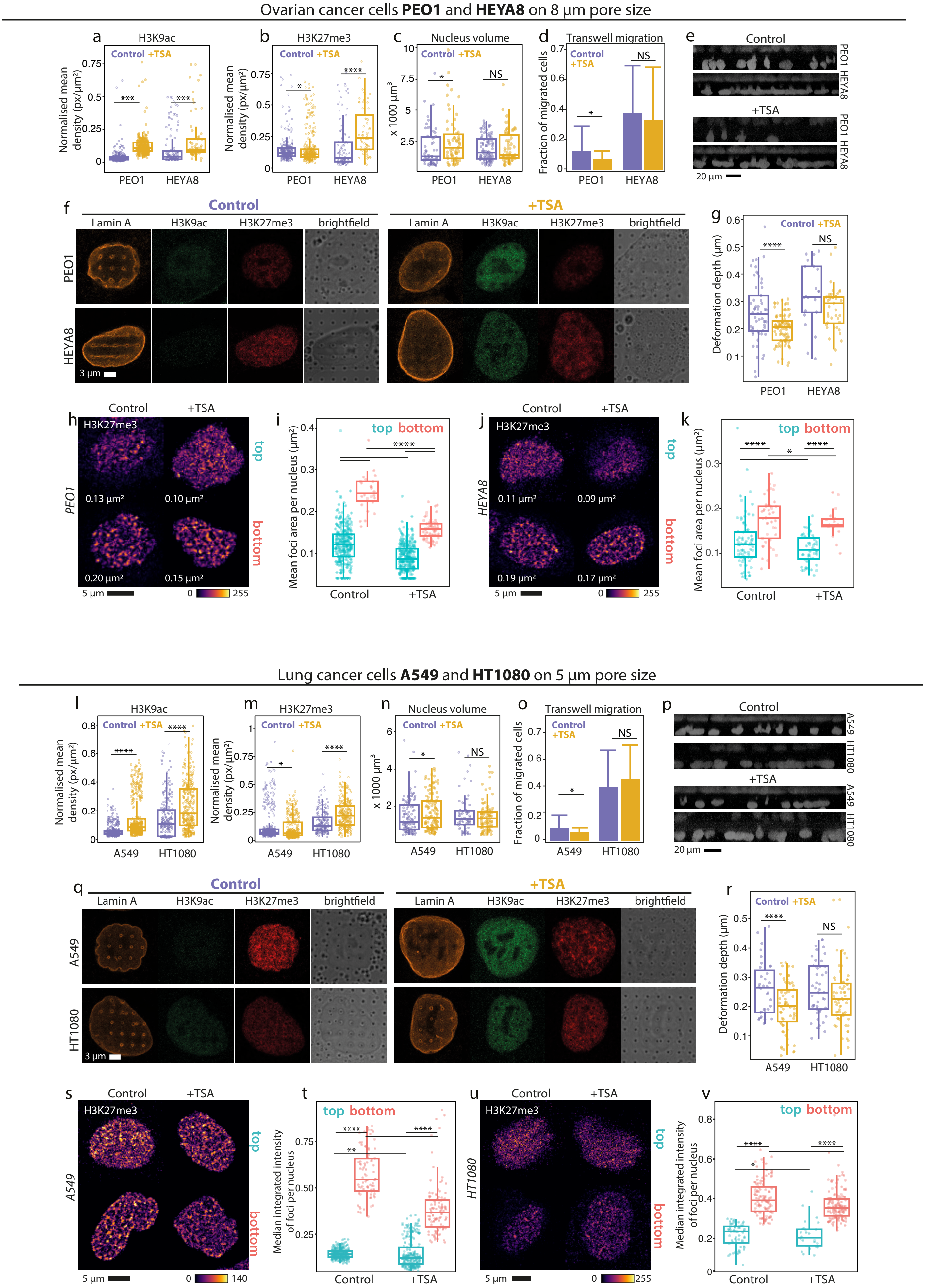
Ovarian cancer cells PEO1 and HEYA8 on 8 μm pore size
a
H3K9ac
Control +TSA
Normalised mean density (px/μm²)
***
***
PEO1
HEYA8
b
H3K27me3
Control +TSA
Normalised mean density (px/μm²)
*
****
PEO1
HEYA8
c
Nucleus volume
Control +TSA
x 1000 μm³
*
NS
PEO1
HEYA8
d
Transwell migration
Control
+TSA
Fraction of migrated cells
*
NS
PEO1
HEYA8
e
Control
PEO1
HEYA8
+TSA
PEO1
HEYA8
20 μm
f
Control
+TSA
Lamin A
H3K9ac
H3K27me3
brightfield
PEO1
HEYA8
3 μm
g
Control +TSA
Deformation depth (μm)
****
NS
PEO1
HEYA8
h
Control
+TSA
H3K27me3
PEO1
top
bottom
0.13 μm²
0.10 μm²
0.20 μm²
0.15 μm²
5 μm
0
255
i
top bottom
Mean foci area per nucleus (μm²)
****
Control
+TSA
j
Control
+TSA
H3K27me3
HEYA8
top
bottom
0.11 μm²
0.09 μm²
0.19 μm²
0.17 μm²
5 μm
0
255
k
top bottom
Mean foci area per nucleus (μm²)
****
*
****
Control
+TSA
Lung cancer cells A549 and HT1080 on 5 μm pore size
l
H3K9ac
Control +TSA
Normalised mean density (px/μm²)
****
****
A549
HT1080
m
H3K27me3
Control +TSA
Normalised mean density (px/μm²)
*
****
A549
HT1080
n
Nucleus volume
Control +TSA
x 1000 μm³
*
NS
A549
HT1080
o
Transwell migration
Control
+TSA
Fraction of migrated cells
*
NS
A549
HT1080
p
Control
A549
HT1080
+TSA
A549
HT1080
20 μm
q
Control
+TSA
Lamin A
H3K9ac
H3K27me3
brightfield
A549
HT1080
3 μm
r
Control +TSA
Deformation depth (μm)
****
NS
A549
HT1080
s
Control
+TSA
H3K27me3
A549
top
bottom
5 μm
0
140
t
top bottom
Median integrated intensity of foci per nucleus
****
****
**
Control
+TSA
u
Control
+TSA
H3K27me3
HT1080
top
bottom
5 μm
0
255
v
top bottom
Median integrated intensity of foci per nucleus
****
*
****
Control
+TSA

Supplementary Figure 17. Highly malignant ovarian and lung cancer cell lines similarly exploit chromatin compaction to counteract acute chromatin decompaction during confined migration.
(a-c) Measures of (a) H3K9ac and (b) H3K27me3 density, and (c) nucleus volume in poorly malignant PEO1 and highly malignant HEYA8 ovarian cancer cell lines.
(d-e) Quantification and representative images of transwell (8 $\mu$m pore size) migration of PEO1 and HEYA8 cells under control and acute TSA-treated conditions.
(f) Representative images of PEO1 and HEYA8 stained against Lamin A, H3K9ac and H3K27me3 nuclei on nanopillars.
(g) Nanopillar-induced deformation depth of PEO1 and HEYA8 under control and acute TSA-treated conditions.
(h-i) Representative images and quantification of size of H3K27me3 foci in PEO1 nuclei after acute TSA treatment.
(j-k) Representative images and quantification of size of H3K27me3 foci in HEYA8 nuclei after acute TSA treatment.
(l-n) Measures of (l) H3K9ac and (m) H3K27me3 density, and (n) nucleus volume in poorly malignant A549 and highly malignant HT1080 lung cancer cell lines.
(d-e) Quantification and representative images of transwell (5 $\mu$m pore size) migration of A549 and HT1080 cells under control and acute TSA-treated conditions.
(f) Representative images of A549 and HT1080 stained against Lamin A, H3K9ac and H3K27me3 nuclei on nanopillars.
(g) Nanopillar-induced deformation depth of A549 and HT1080 under control and acute TSA-treated conditions.
(h-i) Representative images and quantification of size of H3K27me3 foci in A549 nuclei after acute TSA treatment.
(j-k) Representative images and quantification of size of H3K27me3 foci in HT1080 nuclei after acute TSA treatment.
Images of H3K27me3 foci are enhanced for clear visualization of foci. Each dot represents one nucleus with the median of H3K27me3 foci size. Median area of H3K27me3 foci identified from the representative image is included.

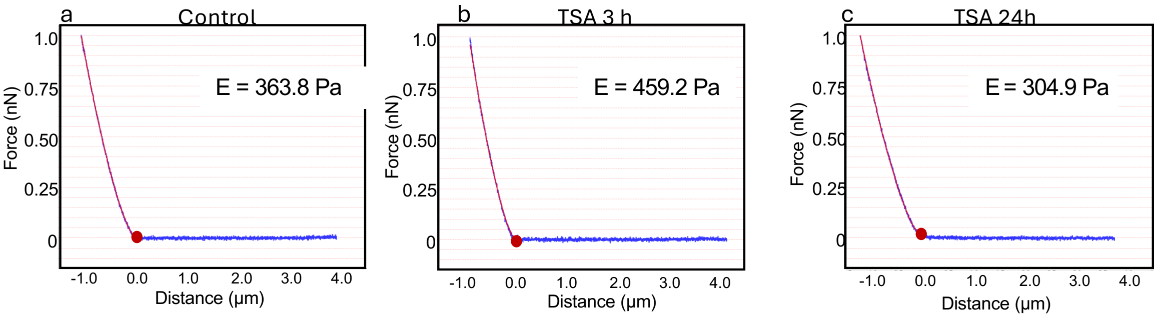


Supplementary Figure 18. Representative force-depth response curves for Hela cells under (a) control, (b) 3-hour TSA and (c) 24-hour TSA treatment.

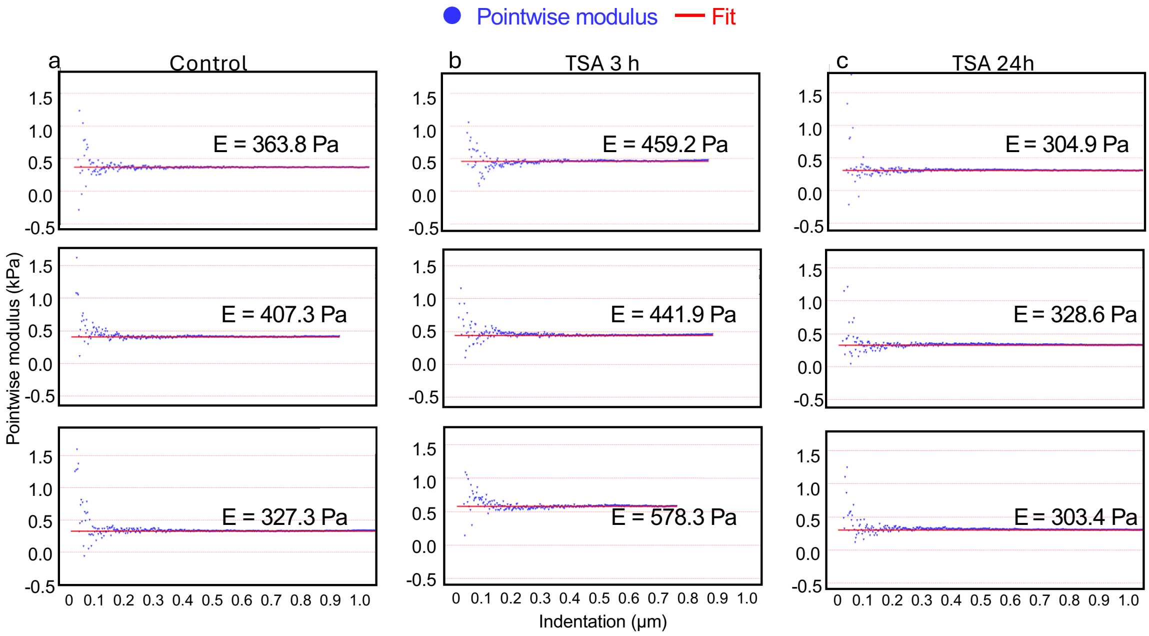


Supplementary Figure 19. Representative pointwise modulus plots for Hela cells under (a) control, (b) 3-hour TSA and (c) 24-hour TSA treatment.
