## Supplementary Info for "Acute chromatin decompaction stiffens the nucleus as revealed by nanopillar-induced nuclear deformation in cells"

[1] School of Chemistry, Chemical Engineering and Biotechnology, Nanyang Technological University, Singapore
[2] Dipartimento di Biologia e Biotecnologie, Sapienza-Università di Roma, Roma, Italy
[3] Department of Biomedical Engineering, National University of Singapore, Singapore
[4] School of Biological Sciences, Nanyang Technological University, Singapore
[5] NTU Institute of Structural Biology (NISB), Experimental Medicine Building, Nanyang Technological University, Singapore
[6] Institute for Health Innovation and Technology (iHealthtech), National University of Singapore, Singapore
[7] Mechanobiology Institute, National University of Singapore, Singapore
[8] Department of Health Science and Technology (D-HEST), ETH Zurich, Zurich, Switzerland
[9] Paul Scherrer Institute, Switzerland
[10] Institute for Digital Molecular Analytics and Science, Nanyang Technological University, Singapore
[#] share co-first authorship

Supplementary Video 1. Hela cell transfected with Lamin A-GFP chromobody showed decreasing deformation depth on nanopillars. Timepoint indicated showed time after TSA treatment.

Supplementary Video 2. Hela cell expressing the diffusive-GFP construct shows change in localization of GFP signal in response to TSA treatment.

### Supplementary methods

*Sample preparation for transmission electron microscopy and data collection:* Hela cells were grown in 35 mm MatTek dishes (MatTek, Ashland, MA, USA, Cat. No. P35G-1.5-14-C) and transfected with 1 µg eGFP-APEX2-H2B plasmid (kindly provided by Eric von Otter; H2B kindly provided by Gabriela Davey, NCBI reference sequence NM_021058.3; for cloning details refer to Hübner et al.[1]) using a 3:1 ratio of PEI (Polysciences, Warrington, PA, USA, Cat. No. 23966; stock: 1 mg/ml) to DNA. The cells were further cultured for 24 h and for TSA treatment 250 ng/mL TSA was added during the last 3.5 h of incubation. Subsequently, cell fixation, sample preparation (based on Lam et al.[2] and Martell et al.[3]) as well as transmission electron microscopy (TEM) data collection was performed according to Hübner et al.[1] Correlation with light microscopy data (eGFP signals and transmission light images) collected during sample preparation was used to identify transfected cells in TEM.

*Quantification of nuclear size and nuclear protein intensities:* Using a customized MATLAB and ImageJ macro codes, nuclei were segmented in 3D by thresholding the DAPI or H3K27me3 fluorescent image across multiple Z planes. The segmented nuclei were used to measure geometrical parameters like nuclear projected area, volume & height as well as fluorescent intensity and density of various proteins in the nucleus. The density of proteins in the nucleus was measured at single cell level by computing the total protein intensity in the nucleus per unit volume or area of the nucleus.

*Calculation of deformation depth induced by nanopillars:* We followed previously published method by Hanson et al.[4] to quantify the deformation depth induced by nanopillars on the nuclear lamina. A custom analysis pipeline was made on Matlab, R and ImageJ. We outline the steps for this analysis below:

1. Image the entire nucleus stained against Lamin A with thinnest z slice possible (0.15-0.3 µm)
2. Using the brightfield z-stack image, select a region of interest centered on the nanopillar of interest. Within this region (1 µm x 1 µm), calculate the mean intensity of

Lamin A at each slice and plot it against z (yielding a z intensity profile). The z intensity profile will contain two distinct maxima. The first maximum denotes the apical of the nucleus (farthest from the nanopillar substrate) while the second maximum denotes the basal side of the nucleus (closer to the nanopillar substrate).

3. To overcome limited resolution in the z axis, fit a polynomial function of order 7 onto the z intensity profile. Using this fitted curve with a continuous z scale (rather than 0.3 µm discrete steps obtained from the z slice intervals), calculate the peak position of the second maximum.
4. Using the brightfield image, select another region of interest surrounding the analysed nanopillar. Calculate the z intensity profile of Lamin A for this region and perform polynomial fitting and calculation of peak position of the second maximum as described in the previous step.
5. Finally, calculate the deformation depth by obtaining the difference in the peak position of the second maxima between the two regions of interest. In the region of interest centered around a nanopillar, the second maximum is closer to the first maximum because the nanopillar indents the nuclear envelope towards the apical side.
6. Calculate the deformation depth of all visible nanopillars that are not at the nucleus periphery and summarise them into a median value that represents the deformability of each nucleus. Multiply this number by the slice interval in µm.

*Generation of intensity profiles of lamins at the nuclear edge:* Confocal images were opened in ImageJ to generate radial intensity profiles of Lamins. ROI of 2 um length were generated from the edge ofthe nucleus and spanning the nuclear envelope, demarcated by either Lamin A or Lamin B1. Using the line profile plug-in in ImageJ, the intensity profile for the Lamin channel was measured across the nuclear envelope. Line profiles were collated and nucleiwere plotted as a scatter plot. The peak of the line profile indicated the enrichment of Lamins at the nuclear envelope with respect to the nucleoplasm.

*Surface cleaning and coating of substrates for expansion microscopy:* Nitrogen air-dried nanopillar substrates were treated with air plasma for 15 minutes, followed by a 20-minute UV treatment. The substrates were then coated with 0.1 mg/ml Poly-L-Lysine (PLL) (Sigma, P4707) for 20 mins, and subsequently washed with PBS, followed by a coating of 0.5% glutaraldehyde (Sigma, G7651) with additional PBS washing steps. For the staining of nanopillars with fluorescently-tagged gelatin, substrates were incubated in gelatin pre-labeled with Atto647N-NHS ester dye (Sigma, 18373) for 1 hour at room temperature, followed by normal cell seeding experiment.

*Expansion microscopy:* After immunostaining, cells were anchored overnight at 4 degrees with a 1:100 dilution of acryloyl-X, SE, (6-((acryloyl)amino) hexanoic acid, succinimidyl ester, Thermo Fisher Scientific, A-20770) in PBS buffer. After anchoring, the sample was washed with PBS. For SA-AA-MBAA based polymerization, we applied the protocol from the Boyden group[5]. The monomer solution, comprise of 19% (w/w) Sodium acrylate (Sigma-Aldrich, 408220), 10% (w/w) acrylamide (Sigma, A8887), 0.1% (w/w) N,N′-Methylenebis(acrylamide) (Sigma, SLBB3006V), was placed on ice and precooled for 5 min. Then 0.5% tetramethyl ethylene diamine (Sigma, 110-18-9) and 0.5% ammonium persulfate (Sigma, A3678) were added to the precooled monomer solution. The mixed solution was then added to the cell and incubate at 4 degrees for 15 minutes, followed by incubating at 37 degrees for 2 h. For the next step, the nanopillar substrates with the hydrogel attached was incubated in digestion buffer with proteinase K (EO0491, Thermo Scientific) and dissolved in the solution comprised of Tris HCl 50 mM, EDTA 1 mM, Triton X-100 0.5%, NaCl 1 M. After 2 hours, the hydrogel was detached from the nanochip. Hydrogels were then immersed in dd$H_2O$ overnight.

*Lattice Light Sheet Imaging:* The hydrogel was transferred to a PLL-coated glass bottom dish, and a few drops of dd$H_2O$ were added to keep the hydrogel humidity during imaging. The Zeiss Lattice Light Sheet 7 microscope was used to image expanded specimens, offering high imaging speed and spatial resolution[6]. The system featured a 44.83x/1 NA water lens for fluorescence signal collection, a 13.3x/0.44 NA lens for illumination, and dual Hamamatsu Orca Fusion sCMOS cameras. Diode lasers at 488 nm (10 mW), 561 nm (10 mW), and 638 nm (5 mW) operated at 30–40% of full power with a 100 ms exposure time. The light sheet had a length of 30 μm along the sample, a thickness of 1000 nm, and a field of view measuring 296.94 μm at 2048 pixels. Post-imaging de-skewing and deconvolution were performed using Zeiss ZEN built-in software with a maximum of 12 iterations of the constrained iterative algorithm.

*Post-imaging calibration in 2 and 3 dimensions:* Local distortion test and calibration was performed using MATLAB and Big Warp plugin with the data analysis methods in previous researchers' study[7]. For the local distortion test, a MATLAB code provided by the Click-ExM group was modified and employed[8].
